# Nanobody CDR3 mimetics enhance SOS1-catalyzed nucleotide exchange on RAS

**DOI:** 10.1101/2022.12.09.519747

**Authors:** Kevin Van holsbeeck, Baptiste Fischer, Simon Gonzalez, Charlène Gadais, Wim Versées, José C. Martins, Charlotte Martin, Alexandre Wohlkönig, Jan Steyaert, Steven Ballet

**Author notes:** Laboratoire de Chimie Biologique, University of Cergy-Pontoise, 5 mail Gay-Lussac, 95031 Cergy-Pontoise, France (Simon Gonzalez). Institut des Sciences Chimiques de Rennes, Equipe CORINT, Université de Rennes 1, 2 Avenue du Pr. Léon Bernard, 35043 Rennes, France. These authors contributed equally (Kevin Van holsbeeck, Baptiste Fischer). All authors have given approval to the final version of the manuscript.

## Abstract

RAS proteins control various intracellular signaling networks. Mutations at specific locations were shown to stabilize their active GTP-bound state, which is associated with the development of multiple cancers. An attractive approach to modulate RAS signaling is through its regulatory guanine nucleotide exchange factor (GEF) son of sevenless 1 (SOS1). With the recent discovery of Nanobody14, which potently enhances SOS1-catalyzed nucleotide exchange on RAS, we explored the feasibility of developing peptide mimetics by structurally mimicking the complementarity-determining region 3 (CDR3). Guided by a biochemical GEF assay and X-ray co-crystal structures, successive rounds of optimization and gradual conformational rigidification led to CDR3 mimetics showing half of the maximal activation potential of the native nanobody. Altogether, this study provides the first proof-of-concept that peptides able to functionally modulate a protein-protein interaction can be obtained by structural mimicry of a nanobody paratope.

---

Antibodies represent an important therapeutic drug family due to their high affinity and specific binding capabilities.^1^ However, they suffer from high production and therapeutic costs, whereas their molecular size (∼150kDa) limits intracellular use. Therefore, downsized functional antibody fragments were developed, such as antigen-binding fragments (Fab, ∼50kDa) and single-chain variable fragments (scFv, ∼25kDa).^2^ Smaller synthetic peptides based on these antibodies’ crucial complementarity-determining regions (CDR) have also been investigated as antibody miniatures, but they frequently show significantly lowered potencies (µM-mM).^3^ Optimization seldomly leads to potent peptidomimetics with biological activities comparable to the native antibody.^4^

The advent and success of nanobodies (Nbs, ∼15kDa) represents a rather untapped alternative source of CDR-based paratopes for the development of low molecular weight (≤2kDa) antibody mimetics.^5^ These small-sized variable heavy chain (VHH) domains maintain the high specificity and efficiency of heterotetrameric antibodies, without needing the significantly larger framework.^6^ Yet, reports of nanobody-based peptidomimetics are limited.^3^ As one of their many interesting applications, nanobodies emerged as useful tools to stabilize conformationally flexible proteins^7^ and protein-protein complexes,^8^ such as the recent discovery of nanobodies modulating the RAS:SOS1 protein complex.^9^

RAS proteins are small G proteins regulating important intracellular signaling networks by functioning as binary molecular switches.^10^ Mutations that lead to a stabilized GTP-bound state can result in aberrant RAS signaling that plays a major role in the development of cancers.^11^ Therefore, the search for direct small molecule RAS inhibitors has received considerable attention, although it turned out to be a major hurdle.^12^ Hence, alternative drug discovery strategies have aimed to address regulators of RAS activation, such as modulation of guanine nucleotide exchange factors (GEFs) like the essential SOS1.^13^ SOS1 catalyzes the conversion of inactive GDP-bound RAS towards its active GTP-bound state. This has inspired the development of several small molecule and peptide inhibitors,^14^ such as small molecules blocking the association of RAS:SOS1,^15^ and α-helix mimetics of SOS1.^16^ Interestingly, antagonist molecules discovered and optimized by Fesik *et al*. reduce the cellular levels of activated RAS (RAS:GTP) upon overactivation.^17^ Similarly to these small molecules, Nanobody14 (Nb14) enhances nucleotide exchange on RAS by interaction with the RAS:SOS1 complex.^9^ Hence, we explored the feasibility to develop downsized peptide mimetics of Nb14 by structurally mimicking one of its CDRs.

The reported RAS:SOS1:Nb14 X-ray structure shows a positioning of Nb14 towards the RAS:SOS1 interface located near the switch II region of RAS (Figure 1A), and reveals a distinct role of the CDRs in the binding event (Figure 1B). CDR1 and CDR2 show a limited number of contacts with SOS1, although CDR1 is positioned over a larger surface on both proteins. In contrast, the top part of CDR3 (Ile^99^-Ala^105^, Figure 1C) makes multiple contacts with SOS1, and is positioned within a hydrophobic pocket in the CDC25 domain of SOS1, which was previously targeted by small molecules.^18^ Consequently, this portion of CDR3 was suggested as an important anchorage point for the binding and activity of Nb14. The tip of this loop adopts a type I β-turn and is supported by two H-bonds between His^100^ and Ser^103^ (Figure 1D). This structural stabilization helps the positioning of the side chains for optimal contact with SOS1, in particular for Trp^102^ of Nb14, which interacts with His^905^, Phe^890^ and Leu^901^ in a hydrophobic binding pocket, and His^100^ with Tyr^884^, His^905^ and Glu^909^ (Figure S1).

**Figure 1.**
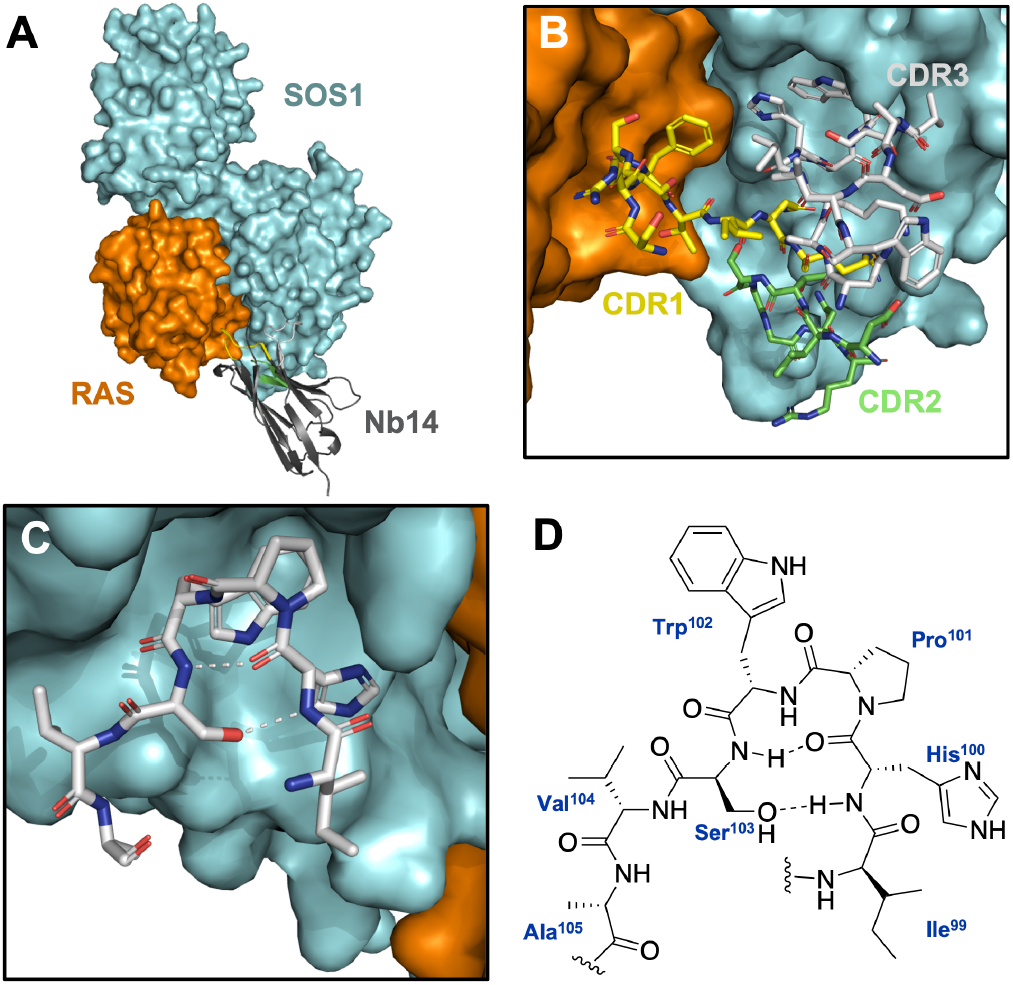
(A) Crystal structure of the RAS:SOS1:Nb14 complex showing the binding interfaces involving Nb14.^9^ (B) Zoom on the CDR loops. (C) Interaction of the truncated CDR3 (Ile99-Ala105) with SOS1. (D) Schematic representation of the truncated CDR3 conformation.

To verify the contribution of the isolated CDR3 on the nucleotide exchange of RAS, linear peptide analogues were synthesized as *C*-terminal amides by standard solid-phase peptide synthesis. Subsequently, the efficacy and potency of these peptides were evaluated via a biochemical nucleotide exchange GEF assay (Figure). As a reference, Nb14 induced a 26-fold increase of SOS1-mediated nucleotide exchange on RAS with an EC_50_ value of 230 nM (Table 1).

**Table 1.** GEF assay results of an initial set of peptide mimetics

| No. | Identity | EC<sub>50</sub> (μM)<sup>a</sup> | Max. Act.<sup>b</sup> |
| --- | --- | --- | --- |
|  | Ref. scaffold R<sup>1</sup> R<sup>2</sup> |  |  |
| Nb14 | / / / | 0.23 ± 0.04 | 25.6 ± 1.2 |
| 1 | A Ac a | 500 ± 140 | 4.3 ± 0.4 |
| 2 | A H b | 166 ± 22 | 8.2 ± 0.3 |
| 3 | B / b | 68 ± 14 | 7.5 ± 0.3 |
| 4 | B / c | 70 ± 15 | 13.8 ± 0.8 |
| 5 | B / d | 36 ± 2 | 8.4 ± 0.1 |

<sup>a</sup>Concentration of the ligands needed to obtain half of the maximum activation rate. <sup>b</sup>Maximum nucleotide exchange rate obtained under saturating ligand concentrations, relative to the activation rate obtained with RAS:SOS1 alone.

As expected, taking CDR3 out of the rigid supporting Nb framework resulted in a significant drop in potency and efficacy. Nevertheless, **1** quadrupled the SOS1-mediated nucleotide exchange on RAS with a low potency (500 µM), which confirmed the importance of this sequence. Therefore, different analogues of **1** were evaluated (Table S1), wherein Pro and Ser were left untouched given their anticipated importance to maintain the β-turn structure. Substitutions of His and Trp mostly led to a loss of activity, which indirectly confirmed their significance for SOS1-interaction.

Although the side chain of Ile^99^ within the CDR3 does not interact with SOS1, it is oriented towards a negatively charged surface patch of SOS1 formed by Glu^902^, Glu^906^, Glu^909^ and Asp^910^ (Figure S1). Hence, substitution of Ile with cationic residues was suggested and proved beneficial for the efficacy and/or potency. In particular, substitution with D-Arg combined with a free *N*-terminus in **2** significantly enhanced the SOS1-mediated nucleotide exchange of RAS, combined with a lower EC_50_ value of 166 µM. To unravel the role of D-Arg in the binding event, a X-ray crystal structure of RAS:SOS1:**2** was determined. In this structure, peptide **2** is located within the same binding site on SOS1 as the Nb14 CDR3 and adopts a similar conformation (Figure S2). The crystal structure reveals that D-Arg does not interact with Glu^906^, Glu^909^ or Asp^910^ as intended, but it forms a H-bond with Tyr^884^ of SOS1 (Figure 3A). Similar to Nb14, Tyr^884^ is H-bonded to His of **2** and forms a cation-π interaction with Arg^73^ of RAS. Noteworthy, His, Pro and Trp of **2** adopt the same orientations as the Nb14 CDR3 (Figure 3B), leading to similar interactions with SOS1. The repositioning of the Ser side chain in **2**, however, does not lead to the formation of the type I β-turn. This observation could be rationalized by the flexible nature of the linear peptidomimetic, which allows the existence of more broadly defined conformer populations in solution.

**Figure 2.**
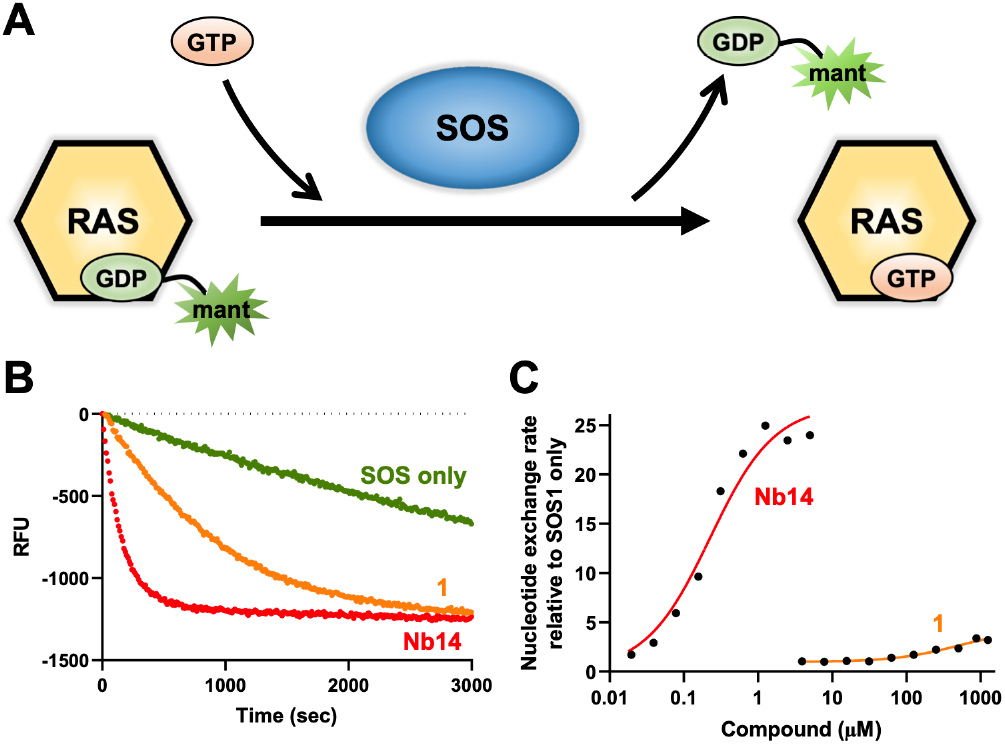
(A) Schematic representation of the GEF assay. The activity of SOS1 is monitored by the decrease of fluorescent signal emitted by labeled GDP (mant-GDP) upon dissociation from RAS. (B) Measured fluorescence in presence of Nb14 (5µM) and **1** (1250µM). (C) Relative SOS1-catalyzed nucleotide exchange rate in presence of Nb14 and **1**.

**Figure 3.**
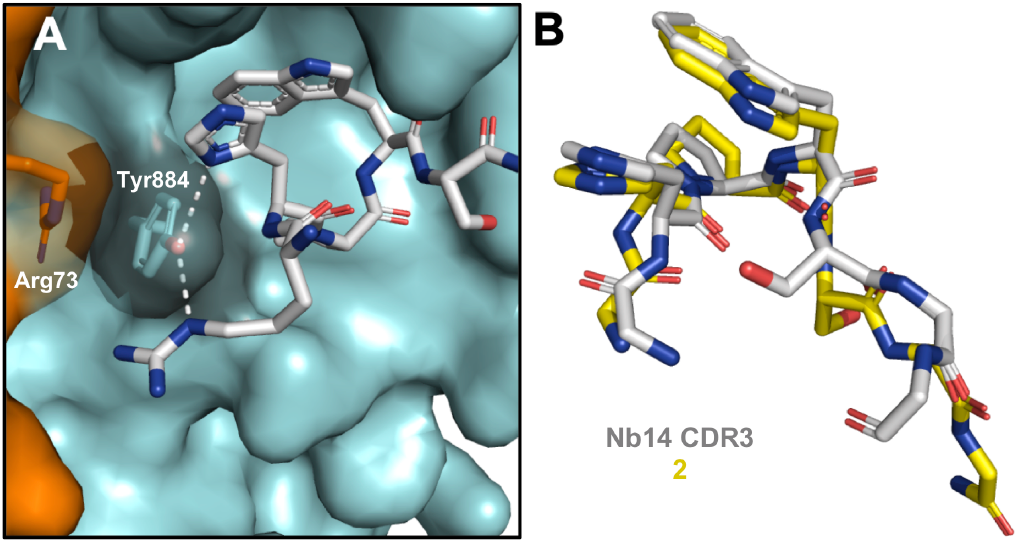
(A) X-ray co-crystal structure of RAS:SOS1:**2**. (B) Superposition of the Nb14 CDR3 and 2. The side chains of residues outside the β-turn were omitted for clarity.

Since the induction of protein-like bioactive conformations by peptide cyclization is well-established,^19^ we considered the use of a lactam bridge between additional Lys and Asp residues. This cyclization strategy resulted in a substantially improved EC_50_ value of 68 µM for **3**, albeit with slightly lower efficacy compared to **2**. The X-ray co-crystal structure of RAS:SOS1:**3** showed a striking difference with mimetic **2** by a reorientation of D-Arg towards Glu^902^ and Glu^906^ of SOS1 (Figure 4A, Figure S3). However, the D-Arg side chain of **3** is not optimally stabilized, being 4-6Å away from Glu^902^ and Glu^906^. Therefore, D-Arg was extended towards D-homoarginine (D-hArg) in **4** and D-Phe(4’-guanidino) in **5**, which led to remarkable improvements. Addition of an extra methylene unit in **4** improved the maximal activation potential to half the value of Nb14. The X-ray structure of RAS:SOS1:**4** (Figure 4B) did reveal the desired extension of D-hArg towards Glu^909^.

**Figure 4.**
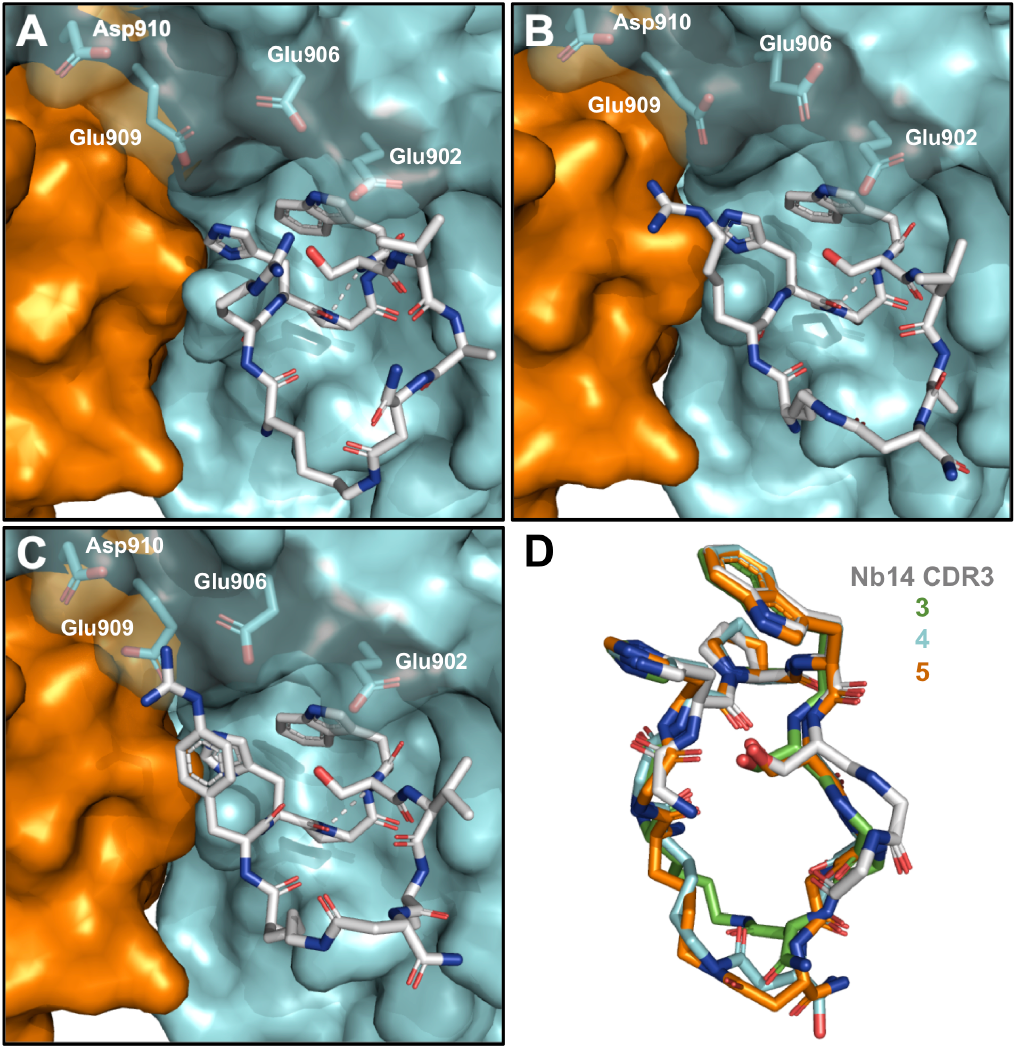
X-ray co-crystal structures of **3** (A), **4** (B) and **5** (C) bound to RAS:SOS1. (D) Superposition of the Nb14 CDR3 and 3-5. The side chains of residues outside the β-turn were omitted for clarity.

In contrast to **4**, mimetic **5** rather improved the EC_50_ value twofold. Here, the X-ray structure of RAS:SOS1:**5** shows an orientation of D-Phe(4’-guanidino) towards Glu^906^ and Glu^909^ (Figure 4C), with both their side chains rotated to optimally interact with the ligand’s guanidyl moiety. The most important interactions explaining the lower EC_50_ value are the salt bridge interaction with Glu^906^, supported by two H-bonds in a bidentate fashion, together with a water-mediated contact with Asp^910^ (Figure S4).

Notably, **3**-**5** show dihedral angles characteristic of the type I β-turn combined with highly preserved orientations of His, Pro, Trp and Ser as compared to the Nb14 CDR3 (Figure 4D), and a H-bond between the carbonyl of His and NH of Ser. However, no productive H-bond is observed between the backbone NH of His and the side chain of Ser. All above observations confirm the structural mimicry of the β-turn found in the Nb14 CDR3 for **3**-**5**, which supports a (partial) functional mimicry. However, residues outside the β-turn region show a larger conformational flexibility, when compared to Nb14.

Given these encouraging results, we proposed that a reduced conformational flexibility by a better induction of the crucial β-turn might reduce the entropic loss upon binding. Since Ala makes no important binding contacts in all X-ray structures, we deleted this residue. Thereby, the crucial β-turn is only flanked by D-Arg and Val, potentially leading to an increased stabilization by the linking bridge. Deletion of Ala in **6** (Table 2) provided a 4-fold improvement of the EC_50_ value compared to **3**, together with a slight activity enhancement. In contrast to the previous series (**4** and **5**, Table 1), the potency improvements of **7** and **8** are rather limited. Additionally, a reduction in the maximal nucleotide exchange rate by **7** was noted, whereas it was improved by **8**.

**Table 1.** GEF assay results of a second set of peptide mimetics

| No. | Identity | EC<sub>50</sub> (μM) | Max. Act. |
| --- | --- | --- | --- |
|  | Ref. scaffold R<sup>2</sup> |  |  |
| **6** | C b | 17.2 ± 2.5 | 8.1 ± 0.2 |
| **7** | C c | 47.5 ± 8.9 | 10.5 ± 0.5 |
| **8** | C d | 28.7 ± 3.1 | 11.5 ± 0.3 |
| **9** | D c | 125 ± 20 | 14.9 ± 0.9 |
| **10** | D d | 87 ± 12 | 14.2 ± 0.6 |

To reduce the conformational freedom at the level of the lactam bridge, the more rigid 1,4-disubstituted 1,2,3-triazole was considered as a linking unit in **9** and **10**. This bridging unit is accessible via a copper(I)-catalyzed azide-alkyne cycloaddition between azidolysine and propargylglycine.^20^ Surprisingly, the GEF assay indicated a loss in potency for both **9** and **10**, although significant efficacy improvements were noted.

**Figure 1.**
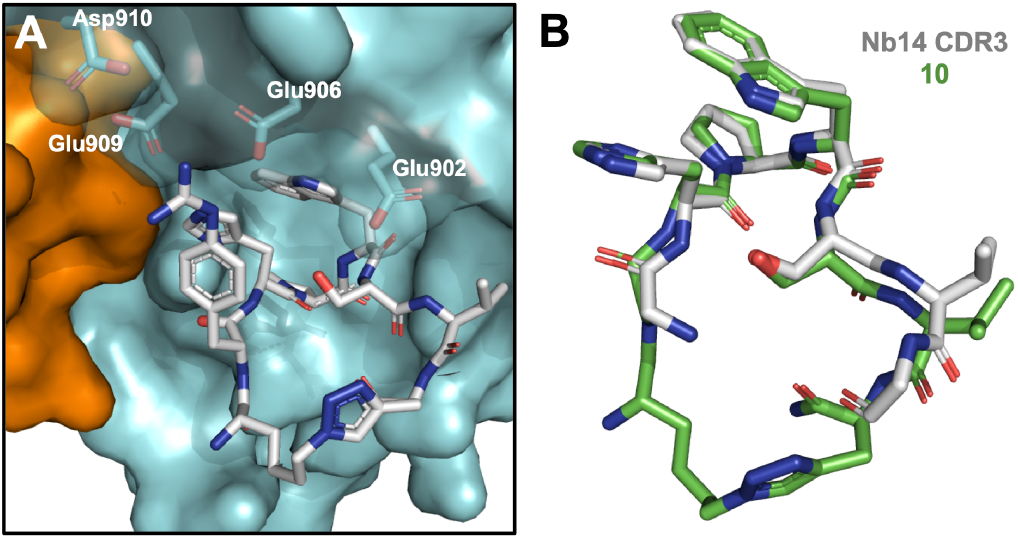
(A) X-ray co-crystal structure of RAS:SOS1:**10**. (B) Superposition of the truncated Nb14 CDR3 and **10**, with omission of certain side chains for clarity.

To rationalize these observations, the X-ray structure of RAS:SOS1:**10** was determined. This structure indicates a similar orientation and binding pattern as **5**, with a rotation of the side chains of Glu^906^ and Glu^909^ allowing the engagement of Glu^906^ by a salt bridge supported by two H-bonds, together with a water-mediated contact with Asp^910^ (Figure 1). In addition, the Lys^899^ side chain is rotated as in **3**-**5**, although the Val side chain of **10** is positioned closer to this residue (Figure S5). Compared to **5**, the Ser side chain is oriented closer towards the His backbone NH, allowing a more productive H-bond as for the Nb14 CDR3 (Figure 1B).

Given the beneficial effect of D-Phe(4’-guanidino) in **8** and **10**, they were studied in more detail. Biolayer interferometry confirmed the binding of biotinylated variants, with affinities of the same order of magnitude as their observed EC_50_ values (respectively 33 and 37 μM, Figure S6). In addition, their effect on the RAS:SOS1 nucleotide exchange catalytic cycle was investigated by measuring all microscopic rate constants in a similar set-up as previously used for Nb14 (Figure S7).^9^ The catalytic cycle is a four step mechanism starting with the formation of the RAS:GDP:SOS1 complex (*K*_*D1*_, Figure 6A), followed by the dissociation of GDP towards the intermediate RAS:SOS1 complex (*K*_*D2*_). The subsequent binding of GTP followed by the accumulation of RAS:GTP, can be considered symmetrical to the first two steps of the catalytic cycle. In the used set-up, Nb14 showed some inhibitory effect on the formation of the RAS:GDP:SOS1 complex (*k*_*1*_ decreased 12-fold, Figure 6B and C), whereas a significant acceleration of the rate limiting transition towards the nucleotide free RAS:SOS1 complex was observed (*k*_*2*_ enhanced 474-fold).^9^ Interestingly, both peptides **8** and **10** displayed a similar effect on the catalytic cycle, although less pronounced (Figures 6B and C). These peptides have a limited effect on the formation of RAS:GDP:SOS1 (*k*_*1*_ decreased 2- to 4-fold), but a major impact on the rate limiting nucleotide release (*k*_*2*_ enhanced 37- to 50-fold). These results indicate that the peptides possess a mode of action comparable to Nb14, which consists of stabilizing a conformation of the RAS:SOS1 complex wherein nucleotides can diffuse in and out with less steric barriers.

Altogether, successive rounds of optimization and gradual conformational rigidification, guided by a biochemical GEF assay and X-ray co-crystal structures, led to CDR3 mimetics showing more than half of the maximal activation potential of the native Nb14. From a biological point of view, these effects are relevant as our best mimetics reach the same activation level generated by the natural allosteric SOS activator.^9, 21^ In addition, a close structural mimicry with the Nb14 CDR3 was observed for mimetic **10**. Thereby, we provided the first proof-of-concept that peptides able to functionally modulate a protein-protein interaction can be obtained by the structural mimicry of a nanobody paratope. Also, the structure-based approach allowed the discovery of functionally important SOS1 residues (Glu^906^, Glu^909^, Asp^910^) which were previously left untouched by small molecule ligands. Subsequent modifications by conformational rigidification and other unnatural amino acids might further improve the potency and activation potential of these peptides.

**Figure 2.**
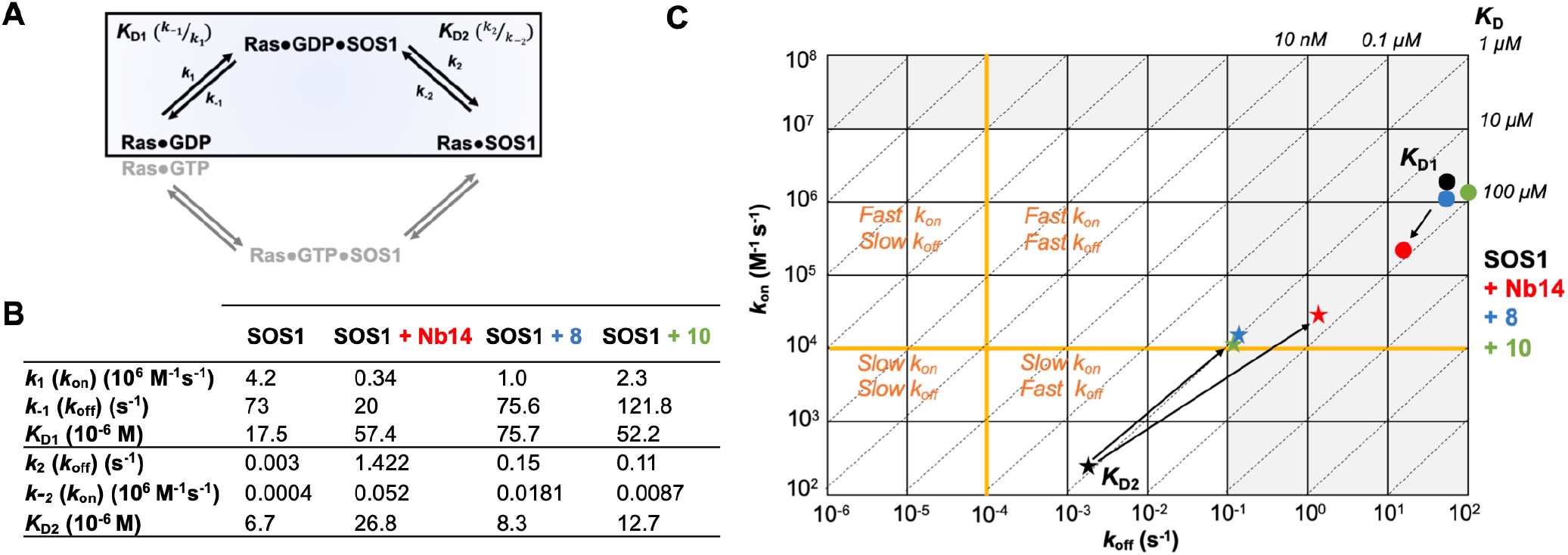
(A) Equilibria involved in the SOS1-catalyzed nucleotide exchange of RAS. (B) Rate and dissociation constants for the interaction between RAS:GDP and SOS1 in presence or absence of Nb14,^9^ **8** and **10**. (C) Kinetic map of SOS1 in presence of Nb14, **8** and **10**. *K*_*D*_ values are shown as diagonal lines, whereas dots and stars indicate the *k*_*on*_ and *k*_*off*_ values obtained for the first (*K*_*D1*_) and second step (*K*_*D2*_) of the catalytic cycle.

## Supporting information

Supplementary figures, tables, procedures and characterisation data

## ASSOCIATED CONTENT

### Supporting Information

The Supporting Information is available free of charge on the ACS Publications website.

Supplementary figures and tables, experimental conditions and characterization data (PDF).

### PDB ID codes

8BE6, 8BE7, 8BE8, 8BE9, 8BEA.

## AUTHOR INFORMATION

### Author Contributions

The manuscript was written through contributions of all authors.

### Notes

The authors declare no competing financial interest.

## ACKNOWLEDGMENT

K.V.h., J.C.M. and S.B thank the Research Foundation Flanders (FWO Vlaanderen) for providing a PhD fellowship to K.V.h. W.V., C.M. and S.B. thank the spearhead (SRP50) program of the Vrije Universiteit Brussel for the financial aid. J.S. and S.B. thank the Research Foundation Flanders for the financial support (S001117N).

---

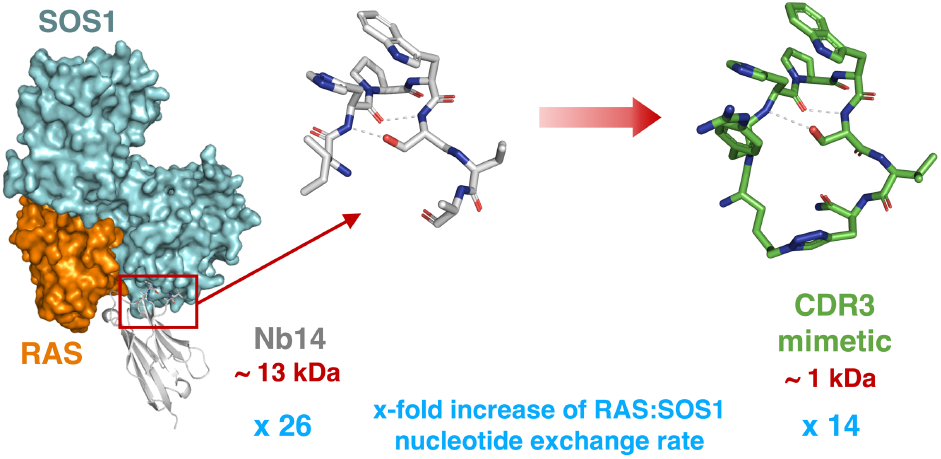

