## Supplementary figures, tables, procedures and characterisation data for "Nanobody CDR3 mimetics enhance SOS1-catalyzed nucleotide exchange on RAS"

[#]Equal contribution

### 1 Supplementary Data

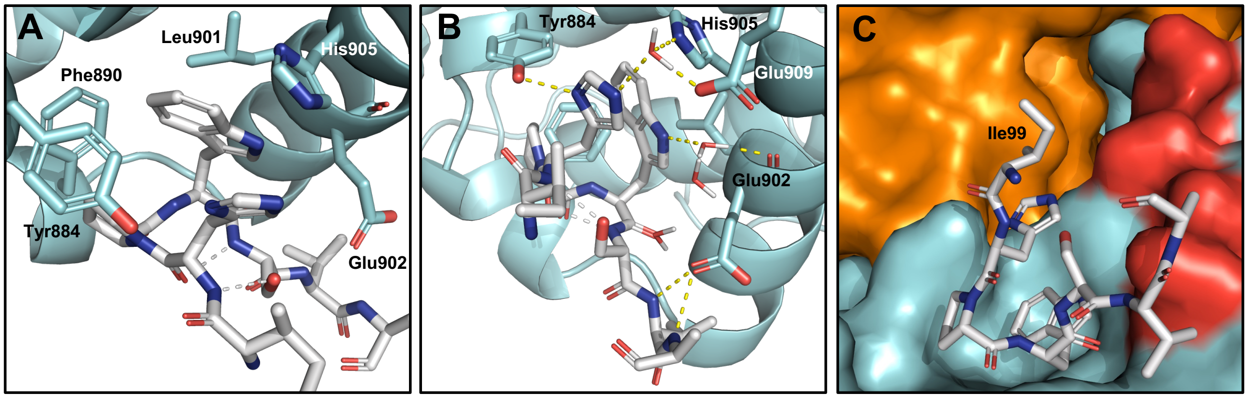

**Figure S1.** Zoom on the interactions of the truncated CDR3 loop of Nb14 with SOS1 found in the RAS:SOS1:Nb14 X-ray structure. $Trp^{102}$ shows hydrophobic contacts with $Phe^{890}$, $Leu^{901}$ and $His^{905}$, and makes a water-mediated contact with the backbone carbonyl of $Glu^{902}$. $His^{100}$ makes a H-bond with $Tyr^{884}$, and water-mediated contacts with $His^{905}$ and $Glu^{909}$. The backbone NH's of $Val^{104}$ and $Ala^{105}$ also engage the carboxylate of $Glu^{902}$ via H-bonds. Panel C shows the orientation of the side chain of $Ile^{99}$ towards a negatively charged surface (colored in red) on SOS1 composed of $Glu^{902}$, $Glu^{906}$, $Glu^{909}$ and $Asp^{910}$.

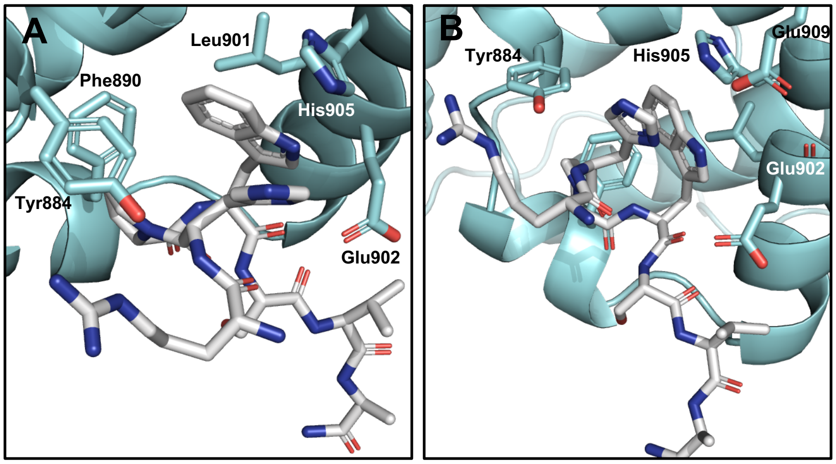

**Figure S2.** Zoom on the interactions of **2** with SOS1 found in the RAS:SOS1:**2** co-crystal X-ray structure.

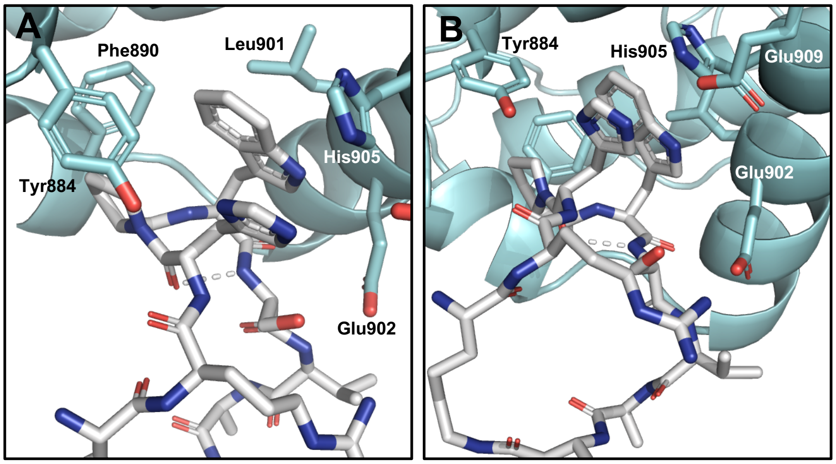

**Figure S3.** Zoom on the interactions of **3** with SOS1 found in the RAS:SOS1:**3** co-crystal X-ray structure.

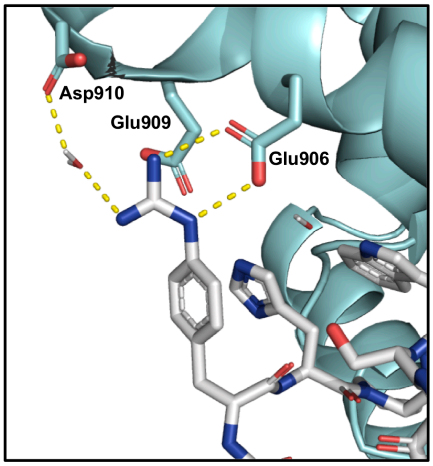

**Figure S4.** Zoom of the RAS:SOS1:**5** co-crystal X-ray structure. The guanidyl function engages $Glu^{906}$ via two H-bonds in a bidentate fashion, together with $Asp^{910}$ via a water-mediated contact.

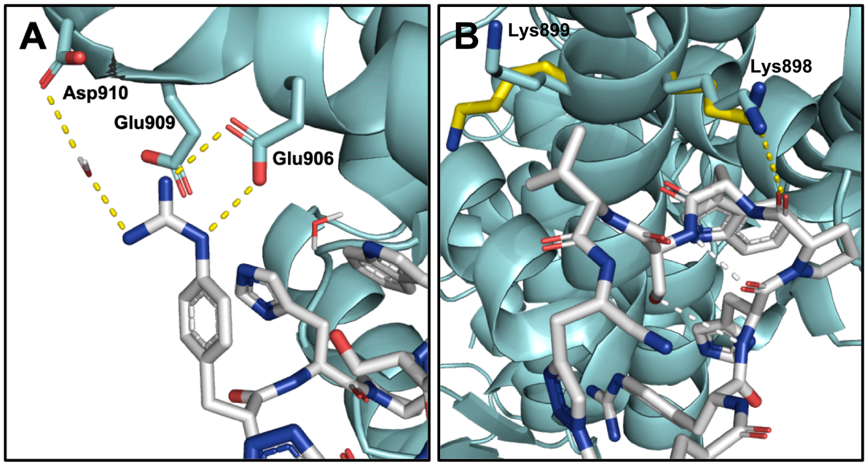

**Figure S5.** Zoom of the RAS:SOS1:**10** co-crystal X-ray structure. (A) The guanidyl function of **10** engages $Glu^{906}$ via two H-bonds in a bidentate fashion, together with $Asp^{910}$ via a water-mediated contact. (B) The Pro carbonyl of **10** contacts $Lys^{898}$, whereas Val is oriented closer towards $Lys^{899}$ compared to **5**. $Lys^{899}$ is rotated compared to the RAS:SOS1:Nb14 X-ray structure (shown in yellow).

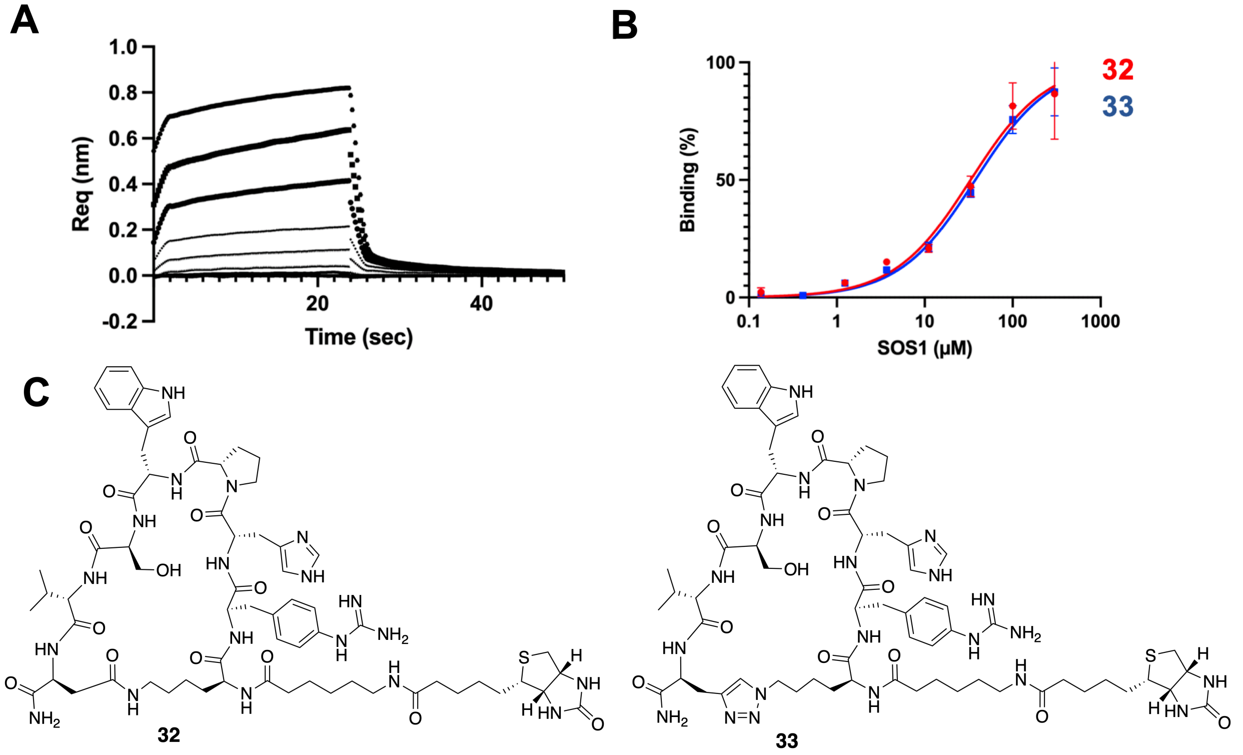

**Figure S6.** Biolayer Interferometry (BLI) measurements to determine the binding affinity of biotinylated variants of **8** and **10** for SOS1 (labelled respectively **32** and **33**). (A) Raw data curves of the association and dissociation steps for **33** and SOS1 determined by BLI. Measurements were performed with biotinylated peptides immobilized on Streptavidin biosensors plunged into solutions containing different concentrations of SOS1 (association step) and then plunged into a buffer solution (dissociation step). (B) Titration curves obtained by plotting normalized signal amplitudes of the association step versus SOS1 concentration. In red, **32** binds SOS1 with an affinity of 33.3 ± 7.3 $\mu$M. In blue, **33** binds SOS1 with an affinity of 37.6 ± 4.7 $\mu$M. (C) Structure of biotinylated peptides **32** and **33**. Aminohexanoic acid (Ahx) was introduced at the *N*-terminus as a spacer between the peptides and biotin.

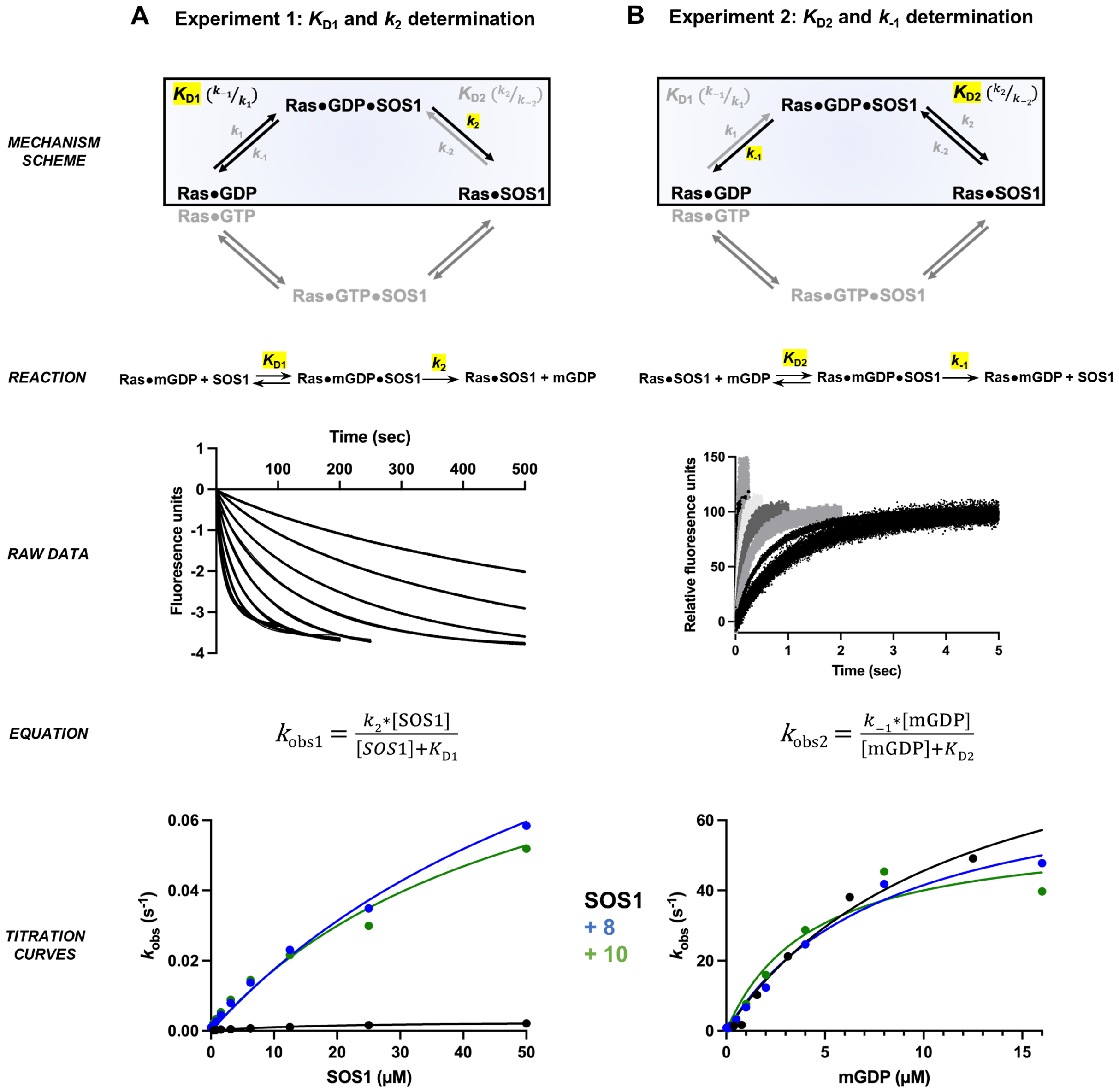

**Figure S7.** Determination of the microscopic rate constants of the SOS1-catalyzed nucleotide exchange reaction of RAS in presence or absence of the peptidomimetics **8** or **10**. Single turnover experiments were performed in a stopped-flow fluorimeter under pseudo first-order conditions to measure all microscopic rate constants in a similar set-up used for Nb14.[1] (A) $K_{D1}$ and $k_2$ were obtained by mixing a fixed concentration (0.5 $\mu$M) of Ras•mGDP with varying excess concentrations of SOS1 and an excess of GDP. Raw data curves were fitted to a single exponential function to determine $k_{obs1}$. Titration curves of $k_{obs1}$ versus SOS1 concentration were plotted and $K_{D1}$ and $k_2$ were determined by deriving $k_{obs1}$ with the indicated equation. A significant increase of $k_{obs}$ and $k_2$ was observed in presence of **8** and **10** (respectively 50- and 37-fold). (B) $K_{D2}$ and $k_{-1}$ were obtained by mixing a fixed concentration (0.5 $\mu$M) of SOS1•Ras complex with varying excess concentrations of mGDP. Raw data curves were fitted to a single exponential equation to determine $k_{obs2}$. Titration curves of $k_{obs2}$ versus SOS1 concentration were plotted and $K_{D2}$ and $k_{-1}$ were determined by deriving $k_{obs2}$ with the indicated equation. Compared to SOS1, no significant change was observed for **8** and **10**.

**Table S1.** GEF assay results of additional synthesized peptidomimetics.

| No. | Sequence | $EC_{50}$ (μM)[a] | Max. Act.[b] |
|---|---|---|---|
| **11** | Ac-Asn-Ala-Lys-Ile-His-Pro-Trp-Ser-Val-Ala-Asp-Leu-Trp-$NH_2$ | 640 ± 90 | 4.1 ± 0.2 |
| **12** | Ac-Ile-**Trp**-Pro-Trp-Ser-Val-Ala-$NH_2$ | ND[c] | ND |
| **13** | Ac-Ile-His-Pro-**D-Trp**-Ser-Val-Ala-$NH_2$ | ND | ND |
| **14** | Ac-Ile-His-Pro-**1-Nal**-Ser-Val-Ala-$NH_2$ | ND | ND |
| **15** | Ac-Ile-His-Pro-**2-Nal**-Ser-Val-Ala-$NH_2$ | ND | ND |
| **16** | Ac-Ile-His-Pro-**Trp(6-Cl)**-Ser-Val-Ala-$NH_2$ | ND | ND |
| **17** | Ac-Ile-His-Pro-**Trp(5-Cl)**-Ser-Val-Ala-$NH_2$ | 2600 ± 2500 | 6 ± 5 |
| **18** | Ac-Ile-His-Pro-**Trp(5,6-diCl)**-Ser-Val-Ala-$NH_2$ | 302 ± 71 | 1.7 ± 0.1 |
| **19** | Ac-**Lys**-His-Pro-Trp-Ser-Val-Ala-$NH_2$ | 1100 ± 180 | 9.5 ± 0.7 |
| **20** | Ac-**Arg**-His-Pro-Trp-Ser-Val-Ala-$NH_2$ | 490 ± 80 | 6.8 ± 0.4 |
| **21** | Ac-**D-Arg**-His-Pro-Trp-Ser-Val-Ala-$NH_2$ | 423 ± 91 | 9.0 ± 0.6 |
| **22** | H-Ile-His-Pro-Trp-Ser-Val-Ala-$NH_2$ | 890 ± 170 | 7.1 ± 0.5 |
| **23** | Ac-**c[**Lys-Ile-His-Pro-Trp-Ser-Val-Ala-Asp**]**-$NH_2$ | 1170 ± 250 | 6.0 ± 0.5 |
| **24** | Ac-**c[**Lys-**Arg**-His-Pro-Trp-Ser-Val-Ala-Asp**]**-$NH_2$ | 880 ± 110 | 12.3 ± 0.7 |
| **25** | H-**c[**Lys-**Arg**-His-Pro-Trp-Ser-Val-Ala-Asp**]**-$NH_2$ | 528 ± 83 | 11.5 ± 0.7 |
| **26** | H-**c[**Lys-**hArg**-His-Pro-Trp-Ser-Val-Ala-Asp**]**-$NH_2$ | 244 ± 44 | 8.2 ± 0.5 |
| **27** | H-Lys-**hArg**-His-Pro-Trp-Ser-Val-Ala-Asp-$NH_2$ | 241 ± 12 | 8.5 ± 0.1 |
| **28** | H-Lys-**D-Arg**-His-Pro-Trp-Ser-Val-Ala-Asp-$NH_2$ | 270 ± 57 | 9.1 ± 0.5 |
| **29** | H-**c**[Lys($N_3$)-**D-Arg**-His-Pro-Trp-Ser-Val-Ala-Pra]-$NH_2$ | 31.0 ± 2.9 | 7.8 ± 0.2 |
| **30** | H-**c**[Orn($N_3$)-**D-Arg**-His-Pro-Trp-Ser-Val-Ala-Pra]-$NH_2$ | 54.3 ± 7.9 | 9.7 ± 0.3 |
| **31** | H-**c**[Dab($N_3$)-**D-Arg**-His-Pro-Trp-Ser-Val-Ala-Pra]-$NH_2$ | 57.0 ± 8.6 | 9.5 ± 0.4 |

[a]$EC_{50}$ values represent the concentration of the peptide needed to obtain half of the maximum activation rate. [b]Max. Act. values represent the maximum activation rate obtained at saturating concentrations of the peptide, and are calculated relative to the activation rate obtained with RAS:SOS1 alone. [c]ND indicates the value was not determined due to an absence of activation measured.

**Table S2.** Data collection and refinement statistics.

| | RAS:SOS1:**2** (8BE6) | RAS:SOS1:**3** (8BE7) | RAS:SOS1:**4** (8BE8) | RAS:SOS1:**5** (8BE9) | RAS:SOS1:**10** (8BEA) |
|---|---|---|---|---|---|
| **Data collection** | | | | | |
| Space group | I422 | I422 | I422 | I422 | I422 |
| Cell dimensions | | | | | |
| *a*, *b*, *c* (Å) | 142.32, 142.32, 207.58 | 143.05, 143.05, 208.33 | 150.06, 150.06, 202.25 | 150.41, 150.41, 200.98 | 150.32, 150.32, 203.71 |
| α, β, γ (°) | 90.00, 90.00, 90.00 | 90.00, 90.00, 90.00 | 90.00, 90.00, 90.00 | 90.00, 90.00, 90.00 | 90.00, 90.00, 90.00 |
| Resolution (Å) | 72.25-2.90 (3.07-2.90) | 47.05-3.00 (3.18-3.00) | 48.57-2.40 (2.54-2.40) | 50-2.51 (2.66-2.51) | 48.66-2.47 (2.56-2.47) |
| $R_{sym}$ or $R_{merge}$ | 0.13 (1.56) | 0.16 (1.86) | 0.14 (1.12) | 0.16 (1.47) | 0.27 (1.49) |
| $I / \sigma I$ | 15.65 (1.61) | 18.88 (1.51) | 19.15 (2.51) | 17.05 (2.03) | 20.93 (2.82) |
| Completeness (%) | 99.0 (99.3) | 99.8 (99.1) | 99.7 (98.6) | 99.2 (95.3) | 99.7 (98.1) |
| Redundancy | 9.75 (9.83) | 26.71 (26.57) | 27.0 (27.2) | 26.2 (26.2) | 27.1 (26.5) |
| **Refinement** | | | | | |
| Resolution (Å) | 46.12-2.90 (3.00-2.90) | 46.48-3.00 (3.11-3.00) | 47.45-2.40 (2.48-2.40) | 48.65-2.51 (2.60-2.51) | 48.66-2.47 (2.56-2.47) |
| No. reflections | 23778 | 21957 | 45398 | 39312 | 41850 |
| $R_{work}$ / $R_{free}$ | 0.225 / 0.270 | 0.209 / 0.260 | 0.199 / 0.231 | 0.1867 / 0.2292 | 0.185 / 0.231 |
| No. atoms | 5071 | 5065 | 5295 | 5291 | 5501 |
| Protein | 4990 | 4982 | 4997 | 5041 | 5066 |
| Ligand/ion | 119 | 77 | 185 | 213 | 146 |
| Water | 20 | 6 | 196 | 124 | 357 |
| *B*-factors | 86.51 | 95.43 | 58.69 | 62.71 | 47.60 |
| Protein | 85.86 | 95.35 | 58.31 | 62.36 | 47.23 |
| Ligand/ion | 144.35 | 128.09 | 83.27 | 80.13 | 65.48 |
| Water | 73.2 | 76.54 | 55.48 | 59.25 | 49.05 |
| R.m.s. deviations | | | | | |
| Bond lengths (Å) | 0.002 | 0.005 | 0.012 | 0.008 | 0.074 |
| Bond angles (°) | 0.49 | 0.95 | 1.11 | 0.89 | 1.48 |

*Values in parentheses are for the highest-resolution shell.

### 2 Chemical synthesis

#### 2.1 General methods

Trifluoroacetic acid (TFA), triisopropylsilane (TIS), phenylsilane, 1-hydroxybenzotriazole hydrate (HOBt·$H_2O$) and copper(I) bromide were purchased from Fluorochem. Fmoc-D-($Boc_2$-4'-guanidino)-Phe-OH originated from Iris Biotech, and Tentagel R RAM resin (0.20 mmol/g) from Rapp Polymere. Other amino acids, coupling reagents and solid supports were purchased from Chem-Impex, whereas all other reagents and solvents were purchased from Sigma Aldrich. Analytical RP-HPLC was carried out on a VWR-Hitachi Chromaster HPLC equipped with a Chromolith High Resolution RP-18C column from Merck (150 mm x 4.6 mm, 1.1 μm; or 50 mm x 4.6 mm, 1.1 μm). Products were detected by a Chromaster HPLC 5430 diode array detector at a wavelength of 214 nm. The solvent system consisted of 0.1% TFA in ultrapure water (A) and 0.1% TFA in acetonitrile (B) with a linear gradient ranging from 1% to 99% B. Preparative RP-HPLC was performed on a Knauer system equipped with a RP-18C ReproSil-Pur ODS-3 column (150 mm x 16 mm, 5 μm), or on a Gilson semi-preparative HPLC equipped with a Supelco Discovery BioWide Pore C18 column (250 mm x 21.2 mm, 10 μm), both equipped with a UV detector set at 214 nm. LC-MS analysis was performed with a Waters 600 HPLC unit equipped with an EC 150/2 NUCLEODUR® 300-5 C18 ec column with a gradient ranging from 3% to 100% of acetonitrile containing 0.1% formic acid, in ultrapure water containing 0.1% formic acid. Monitoring was done via UV detection at 214 nm. Coupled MS analysis was performed on a Micromass QTOF-micro system. HRMS was conducted on the same device with reserpine as the reference.

#### 2.2 General peptide synthesis methods

##### *Peptide couplings*

Peptides were synthesized on a Rink Amide AM resin (loading 0.33-0.47 mmol/g) on 0.05 to 0.15 mmol scales by iterative cycles of $N^{\alpha}$-Fmoc deprotection, amino acid couplings, and washing steps, either manually or in an automated fashion using a Liberty Lite™ Automated Microwave Peptide Synthesizer. Amino acids were used in their $N^{\alpha}$-Fmoc-protected form, combined with the following side chain protecting groups: trityl (Trt) for Asn and His; *tert*-butyl (*t*Bu) for Ser, Thr and Asp; *tert*-butyloxycarbonyl (Boc) for Lys, Trp and D-Trp; 2,2,4,6,7-pentamethyldihydrobenzofuran-5-sulfonyl (Pbf) for Arg, D-Arg, hArg and D-hArg; and di-*tert*-butyloxycarbonyl ($Boc_2$) for D-Phe-(4'-guanidino).

On the Liberty Lite™ peptide synthesizer, couplings were performed using 4 equiv. of amino acids (0.5 M solution in DMF), *N,N'*-diisopropylcarbodiimide (DIC, 0.5 M solution in DMF), and Oxyma Pure (1 M solution in DMF). Whereas standard couplings were performed for 2.1 minutes at 90°C, coupling of Fmoc-His(Trt)-OH was performed for 10 minutes at 50°C, and Fmoc-Arg(Pbf)-OH was coupled twice at 75°C. Fmoc deprotections were carried out for 1.05 minutes at 90°C using 20 % (v/v) 4-methylpiperidine in DMF.

Manual syntheses were carried out in polypropylene syringes equipped with a polyethylene frit. Between every step, the resin was washed thoroughly with DMF (3 times) and DCM (3 times). Fmoc deprotections were performed using 20 % (v/v) 4-methylpiperidine in DMF for 5 and 15 minutes. Standard amino acid couplings were carried out using 3 equiv. of protected amino acid, 3 equiv. of *N,N,N',N'*-tetramethyl-O-(1H-benzotriazol-1-yl)uronium hexafluorophosphate

(HBTU) and 5 equiv. of *N*,*N*-diisopropylethylamine (DIPEA) in DMF for 45 to 90 minutes at room temperature. *N*-terminal acetylation was carried out using acetic anhydride (10 equiv.) and DIPEA (10 equiv.) in DCM for 30 minutes. Completion of couplings was verified at all stages by Kaiser or chloranil colorimetric tests and/or small-scale cleavages followed by LC-MS analyses.

***Peptide cleavages and purification***

Peptide cleavages were performed with a freshly prepared solution of TFA/TIS/$H_2O$ 95/2.5/2.5 (v/v/v) for 2 h at room temperature, with an extension towards 4 h for sequences containing a Pbf protecting group. Subsequently, the resin was filtered and washed with DCM and neat TFA. The combined filtrates were evaporated. The crude peptides were obtained after precipitation in cold diethyl ether, centrifugation and lyophilization. Purifications were subsequently performed by preparative RP-HPLC using a linear gradient, yielding peptides with a HPLC purity over 95%.

#### 2.3 Synthesis of peptidomimetics

***Synthesis of linear peptidomimetics 1, 11-13, 19-20, 22***

Peptides were synthesized using the Liberty Lite™ peptide synthesizer, whereas *N*-terminal acetylations and final cleavages were performed manually as outlined above.

***Synthesis of linear peptidomimetics 2, 14-18, 21***

Peptides were synthesized manually as outlined above. Non-standard residues (1-Nal, 2-Nal, Trp(6-Cl), Trp(5-Cl), Trp(5,6-diCl), D-Arg) were coupled using 1.5 equiv. of protected amino acid, 1.5 equiv. of HBTU and 2.5 equiv. of DIPEA in DMF for 2 h.

***Synthesis of lactam cyclized mimetics 3-8, 23-28***

Peptides were initially synthesized in a linear fashion, wherein *N*-Fmoc protected Asp and Lys were used, respectively, under their allyl (Allyl) and allyloxycarbonyl (Alloc) side chain protected forms. Amino acid residues Ile, His, Pro, Trp, Ser, Val and Ala were attached to the resin using the Liberty Lite™ peptide synthesizer, whereas all other residues were coupled manually using 1.5 equiv. of protected amino acid, 1.5 equiv. of HBTU and 2.5 equiv. of DIPEA in DMF for 90 minutes. Subsequently, the Allyl and Alloc protecting groups were removed by treatment with a solution of tetrakis(triphenylphosphine)palladium (0.25 equiv.) and phenylsilane (24 equiv.) in DCM for two times 30 minutes at room temperature. After thorough washing of the resin with DMF (2 times), a 11.6 mM solution of diethyldithiocarbamate in DMF containing 0.01 % (v/v) DIPEA (6 times), DMF (2 times) and DCM (3 times), the resin-bound peptides were cyclized overnight using HBTU (6 equiv.), HOBt·$H_2O$ (6 equiv) and DIPEA (12 equiv.) in DMF at room temperature. Completion of the couplings was verified by Kaiser tests and/or small-scale cleavages followed by LC-MS analysis. Final Fmoc removal and subsequent steps were identical as outlined before.

***Synthesis of CuAAC cyclized peptidomimetics 9-10***

Peptides were synthesized in a linear fashion on a Tentagel R RAM resin (loading 0.20 mmol/g) using an identical protocol as outlined above. Amino acid residues His, Pro, Trp, Ser and Val were coupled using the Liberty Lite™ peptide synthesizer, whereas all other residues were coupled manually using 1.5 equiv. of protected amino acid, 1.5 equiv. of HBTU and 3.0 equiv. of DIPEA in DMF for 90 minutes. Cyclization was performed before final Fmoc removal using slightly modified conditions of a previously published procedure.[2] After swelling of the resin in DCM for 10 minutes, copper(I)bromide (3 equiv.) completely dissolved in degassed DMSO (55 ml/mmol) was added, followed by sodium ascorbate (3 equiv.) dissolved in water (10 ml/mmol), 2,6-lutidine (10 equiv.) and DIPEA (10 equiv.). After bubbling Ar through the mixture for 5 minutes, the syringe was closed and shaken for 24 h at room temperature. Subsequently, the resin was washed with 2-propanol/DMSO 5/3 (v/v, 2 times), a 1.0 M solution of pyridine·HCl in DCM/MeOH 95/5 (v/v, 3 times), DMF (3 times) and DCM (3 times). After final Fmoc removal, the peptides were cleaved and purified as outline before.

***Synthesis of CuAAC cyclized mimetics 29-31***

Peptides were synthesized manually in a linear fashion, using the protocol outlined above. Non-standard residues (Pra, D-Arg, Lys($N_3$), Orn($N_3$), Dab($N_3$)) were coupled manually using 1.5 equiv. of protected amino acid, 1.5 equiv. of HBTU and 2.5 equiv. of DIPEA in DMF for 90 minutes. After final Fmoc removal, the resin was swollen in DCM, and washed twice with DMF degassed with Ar. Subsequently, copper(I)bromide (3 equiv.) dissolved in degassed DMF (40 ml/mmol) was added, followed by sodium ascorbate (3 equiv.) dissolved in water (3 ml/mmol), 2,6-lutidine (10 equiv.) and DIPEA (10 equiv.). After bubbling Ar through the mixture for 5 minutes, the syringe was sealed and shaken for 18 h at room temperature. Subsequently, the resin was washed with DMF (2 times), a 1.0 M solution of pyridine·HCl in DCM/MeOH 95/5 (v/v, 3 times), DMF (3 times), 2-propanol and DCM (3 times). Successful cyclization was verified by performing a small-scale cleavage followed by HPLC-MS analyses, which indicated a shift in retention times compared to the linear starting materials. Subsequent steps were identical as outlined before.

***Synthesis of biotinylated peptides 32***

Peptide **32** was synthesized in an identical manner as peptide **8**. However, after final Fmoc deprotection, the peptide was further elongated with Fmoc-Ahx-OH (2.0 equiv.) using HBTU (2.0 equiv.) and DIPEA (4.0 equiv.) in DMF for 2 hours. After subsequent Fmoc deprotection, biotin (4.0 equiv.) was coupled using HBTU (4.0 equiv.) an DIPEA (8.0 equiv.) in DMSO/DMF 1/1 for 5 hours. After washes with DMSO (3 times), DMF (3 times) and DCM (3 times), the peptides were cleaved and purified as outlined above.

***Synthesis of biotinylated peptides 33***

Peptide **33** was synthesized in an identical manner as peptide **10**. However, after final Fmoc deprotection, the peptide was further elongated with Fmoc-Ahx-OH (2.0 equiv.) using HBTU (2.0 equiv.) and DIPEA (4.0 equiv.) in DMF for 2 hours. After subsequent Fmoc deprotection, biotin (4.0 equiv.) was coupled using HBTU (4.0 equiv.) an DIPEA (8.0 equiv.) in DMSO/DMF 1/1 for 5 hours. After washes with DMSO (3 times), DMF (3 times) and DCM (3 times), the peptides were cleaved and purified as outlined above.

#### 2.4 Analytical characterization of synthesized peptides

**Table S3.** Characterization of synthesized peptides (part 1).

| No. | Sequence | Molecular formula | HPLC | | | HRMS $[M+X]^+$ | | |
|---|---|---|---|---|---|---|---|---|
| | | | Method* | r.t. (min) | Purity | X | Calculated | Found |
| **1** | Ac-Ile-His-Pro-Trp-Ser-Val-Ala-NH2 | $C_{41}H_{59}N_{11}O_9$ | 1 | 2.88 | > 98% | H | 850.4575 | 850.4540 |
| **2** | H-**D-Arg**-His-Pro-Trp-Ser-Val-Ala-NH2 | $C_{39}H_{58}N_{14}O_8$ | 2 | 1.60 | > 98% | H | 851.4641 | 851.4631 |
| **3** | H-**c**[Lys-**D-Arg**-His-Pro-Trp-Ser-Val-Ala-Asp]-NH2 | $C_{49}H_{74}N_{17}O_{11}$ | 2 | 1.64 | > 97% | H | 1076.5753 | 1076.5730 |
| **4** | H-**c**[Lys-**D-hArg**-His-Pro-Trp-Ser-Val-Ala-Asp]-NH2 | $C_{50}H_{75}N_{17}O_{11}$ | 1 | 2.59 | > 98% | H | 1090.5911 | 1090.5935 |
| **5** | H-**c**[Lys-**D-Phe(4'-guanidino)**-His-Pro-Trp-Ser-Val-Ala-Asp]-NH2 | $C_{53}H_{73}N_{17}O_{11}$ | 1 | 2.60 | > 98% | H | 1124.5753 | 1124.5682 |
| **6** | H-**c**[Lys-**D-Arg**-His-Pro-Trp-Ser-Val-Asp]-NH2 | $C_{46}H_{68}N_{16}O_{10}$ | 1 | 3.13 | > 96% | H | 1005.5383 | 1005.5405 |
| **7** | H-**c**[Lys-**D-hArg**-His-Pro-Trp-Ser-Val-Asp]-NH2 | $C_{47}H_{71}N_{16}O_{10}$ | 1 | 2.55 | > 98% | H | 1019.5539 | 1019.5479 |
| **8** | H-**c**[Lys-**D-Phe(4'-guanidino)**-His-Pro-Trp-Ser-Val-Asp]-NH2 | $C_{50}H_{68}N_{16}O_{10}$ | 1 | 2.55 | > 96% | H | 1053.5382 | 1053.5431 |
| **9** | H-**c**[Lys(N3)-**D-hArg**-His-Pro-Trp-Ser-Val-Pra]-NH2 | $C_{51}H_{70}N_{18}O_9$ | 1 | 2.53 | > 98% | H | 1043.5652 | 1043.5651 |
| **10** | H-**c**[Lys(N3)-**D-Phe(4'-guanidino)**-His-Pro-Trp-Ser-Val-Pra]-NH2 | $C_{48}H_{69}N_{18}O_9$ | 1 | 2.54 | > 98% | H | 1077.5494 | 1077.5404 |

*HPLC method 1: Performed with a Chromolith High Resolution RP-18C column (150 mm x 4.6 mm, 1.1 µm) using a linear gradient ranging from 1% to 99% of acetonitrile containing 0.1% TFA in ultrapure water containing 0.1% TFA, over 6 minutes.

HPLC method 2: Performed with a Chromolith High Resolution RP-18C column (50 mm x 4.6 mm, 1.1 µm) using a linear gradient ranging from 1% to 99% of acetonitrile containing 0.1% TFA in ultrapure water containing 0.1% TFA, over 5.5 minutes.

**Table S4.** Characterization of synthesized peptides (part 2).

| No. | Sequence | Molecular formula | HPLC | | | HRMS $[M+X]^+$ | | |
|---|---|---|---|---|---|---|---|---|
| | | | Method | r.t. (min) | Purity | X | Calculated | Found |
| **11** | Ac-Asn-Ala-Lys-Ile-His-Pro-Trp-Ser-Val-Ala-Asp-Leu-Trp-NH$_2$ | $C_{75}H_{108}N_{20}O_{18}$ | 1 | 3.19 | > 98% | H | 1577.8229 | 1577.8171 |
| **12** | Ac-Ile-**Trp**-Pro-Trp-Ser-Val-Ala-NH$_2$ | $C_{46}H_{62}N_{10}O_9$ | 2 | 2.31 | > 97% | Na | 921.4599 | 921.4576 |
| **13** | Ac-Ile-His-Pro-**D-Trp**-Ser-Val-Ala-NH$_2$ | $C_{41}H_{59}N_{11}O_9$ | 1 | 2.96 | > 98% | H | 850.4575 | 850.4586 |
| **14** | Ac-Ile-His-Pro-**1-Nal**-Ser-Val-Ala-NH$_2$ | $C_{43}H_{60}N_{10}O_9$ | 2 | 2.11 | > 98% | H | 861.4623 | 861.4694 |
| **15** | Ac-Ile-His-Pro-**2-Nal**-Ser-Val-Ala-NH$_2$ | $C_{43}H_{60}N_{10}O_9$ | 1 | 3.13 | > 98% | Na | 883.4442 | 883.4435 |
| **16** | Ac-Ile-His-Pro-**Trp(6-Cl)**-Ser-Val-Ala-NH$_2$ | $C_{41}H_{58}ClN_{11}O_9$ | 2 | 2.06 | > 98% | H | 884.4186 | 884.4183 |
| **17** | Ac-Ile-His-Pro-**Trp(5-Cl)**-Ser-Val-Ala-NH$_2$ | $C_{41}H_{58}ClN_{11}O_9$ | 2 | 2.06 | > 98% | Na | 906.4005 | 906.3983 |
| **18** | Ac-Ile-His-Pro-**Trp(5,6-diCl)**-Ser-Val-Ala-NH$_2$ | $C_{41}H_{57}Cl_2N_{11}O_9$ | 2 | 2.17 | > 98% | H | 918.3796 | 918.3710 |
| **19** | Ac-**Lys**-His-Pro-Trp-Ser-Val-Ala-NH$_2$ | $C_{41}H_{60}N_{12}O_9$ | 2 | 1.61 | > 98% | H | 865.4684 | 865.4636 |
| **20** | Ac-**Arg**-His-Pro-Trp-Ser-Val-Ala-NH$_2$ | $C_{41}H_{60}N_{14}O_9$ | 2 | 1.65 | > 97% | H | 893.4746 | 893.4694 |
| **21** | Ac-**D-Arg**-His-Pro-Trp-Ser-Val-Ala-NH$_2$ | $C_{41}H_{60}N_{14}O_9$ | 1 | 2.52 | > 96% | H | 893.4746 | 893.4757 |
| **22** | H-Ile-His-Pro-Trp-Ser-Val-Ala-NH$_2$ | $C_{39}H_{57}N_{11}O_8$ | 2 | 1.73 | > 98% | Na | 830.4290 | 830.4213 |
| **23** | Ac-**c**[Lys-Ile-His-Pro-Trp-Ser-Val-Ala-Asp]-NH$_2$ | $C_{51}H_{74}N_{14}O_{12}$ | 2 | 1.88 | > 98% | H | 1075.5688 | 1075.5624 |
| **24** | Ac-**c[**Lys-**Arg**-His-Pro-Trp-Ser-Val-Ala-Asp**]**-NH$_2$ | $C_{51}H_{75}N_{17}O_{12}$ | 1 | 2.61 | > 98% | H | 1118.5859 | 1118.5852 |
| **25** | H-**c[**Lys-**Arg**-His-Pro-Trp-Ser-Val-Ala-Asp**]**-NH$_2$ | $C_{49}H_{73}N_{17}O_{11}$ | 1 | 2.53 | > 98% | H | 1076.5753 | 1076.5769 |
| **26** | H-**c[**Lys-**hArg**-His-Pro-Trp-Ser-Val-Ala-Asp**]**-NH$_2$ | $C_{50}H_{75}N_{17}O_{11}$ | 2 | 1.68 | > 97% | H | 1090.5911 | 1090.5884 |
| **27** | H-Lys-**hArg**-His-Pro-Trp-Ser-Val-Ala-Asp-NH$_2$ | $C_{50}H_{77}N_{17}O_{12}$ | 2 | 1.57 | > 98% | Na | 1130.5835 | 1130.5854 |
| **28** | H-Lys-**D-Arg**-His-Pro-Trp-Ser-Val-Ala-Asp-NH$_2$ | $C_{49}H_{75}N_{17}O_{12}$ | 2 | 1.57 | > 98% | H | 1094.5859 | 1094.5831 |
| **29** | H-**c**[Lys(N$_3$)-**D-Arg**-His-Pro-Trp-Ser-Val-Ala-Pra]-NH$_2$ | $C_{50}H_{73}N_{19}O_{10}$ | 1 | 2.56 | > 98% | H | 1100.5865 | 1100.5916 |
| **30** | H-**c**[Orn(N$_3$)-**D-Arg**-His-Pro-Trp-Ser-Val-Ala-Pra]-NH$_2$ | $C_{49}H_{71}N_{19}O_{10}$ | 1 | 2.51 | > 98% | H | 1086.5709 | 1086.5692 |
| **31** | H-**c**[Dab(N$_3$)-**D-Arg**-His-Pro-Trp-Ser-Val-Ala-Pra]-NH$_2$ | $C_{48}H_{69}N_{19}O_{10}$ | 1 | 2.58 | > 98% | H | 1072.5553 | 1072.5533 |
| **32** | Biotin-Ahx-**c**[Lys-**D-Phe(4'-guanidino)**-His-Pro-Trp-Ser-Val-Asp]-NH$_2$ | $C_{66}H_{93}N_{19}O_{13}S$ | 1 | 2.77 | > 99% | Na | 1414.6819 | 1414.6812 |
| **33** | Biotin-Ahx-**c**[Lys(N$_3$)-**D-Phe(4'-guanidino)**-His-Pro-Trp-Ser-Val-Pra]-NH$_2$ | $C_{67}H_{93}N_{21}O_{12}S$ | 1 | 2.79 | > 97% | H | 1416.7112 | 1416.7081 |

#### 2.5 HPLC chromatograms of synthesized peptides

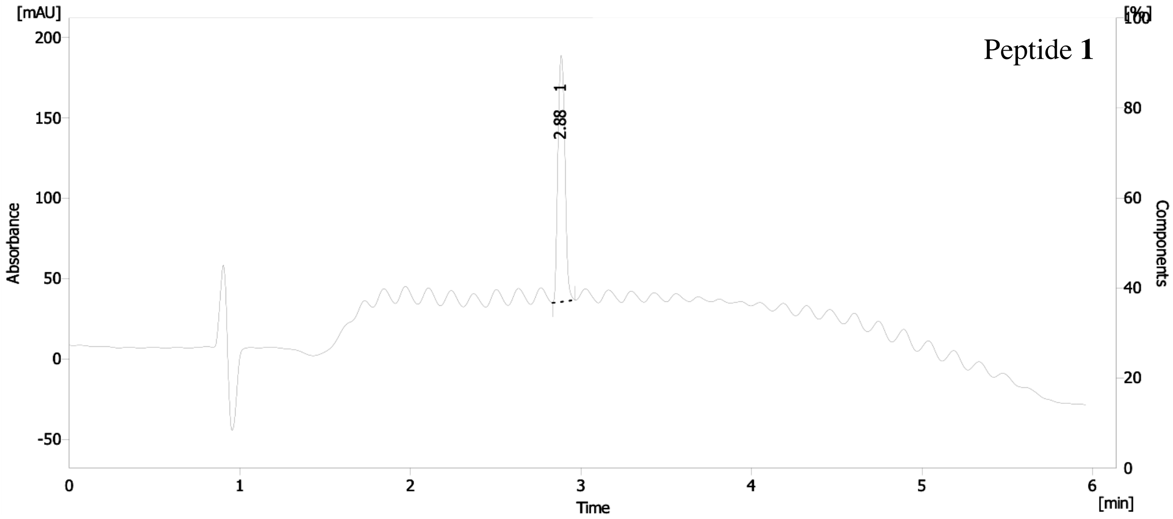

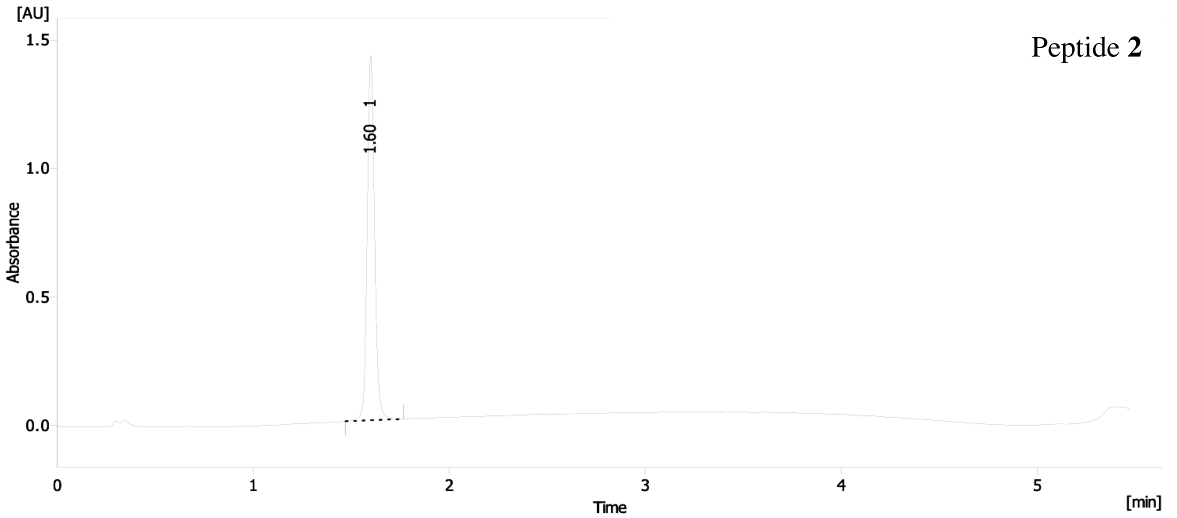

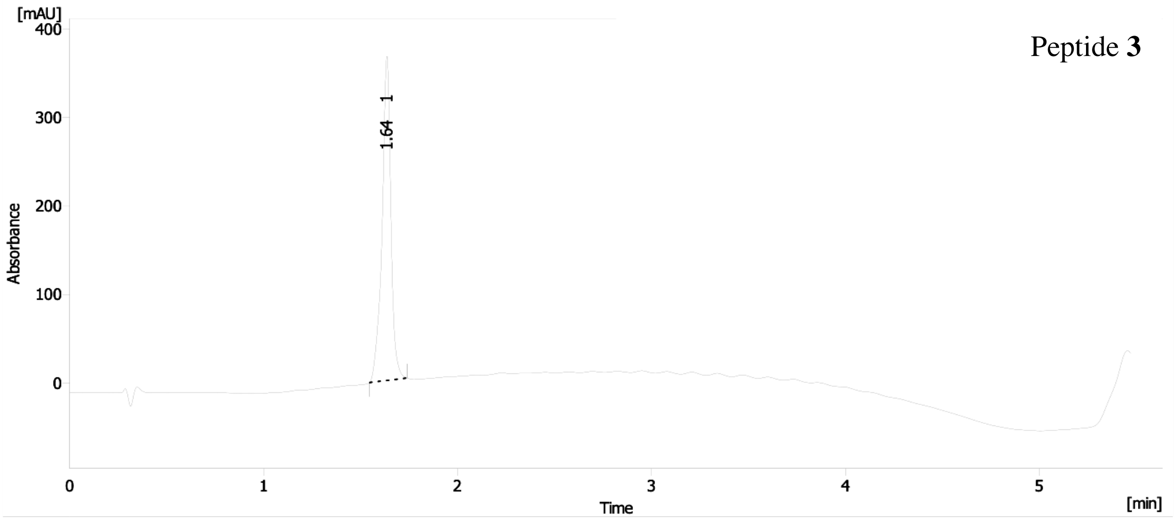
Peptide 3
[mAU]
Absorbance
1.64 1
Time
[min]

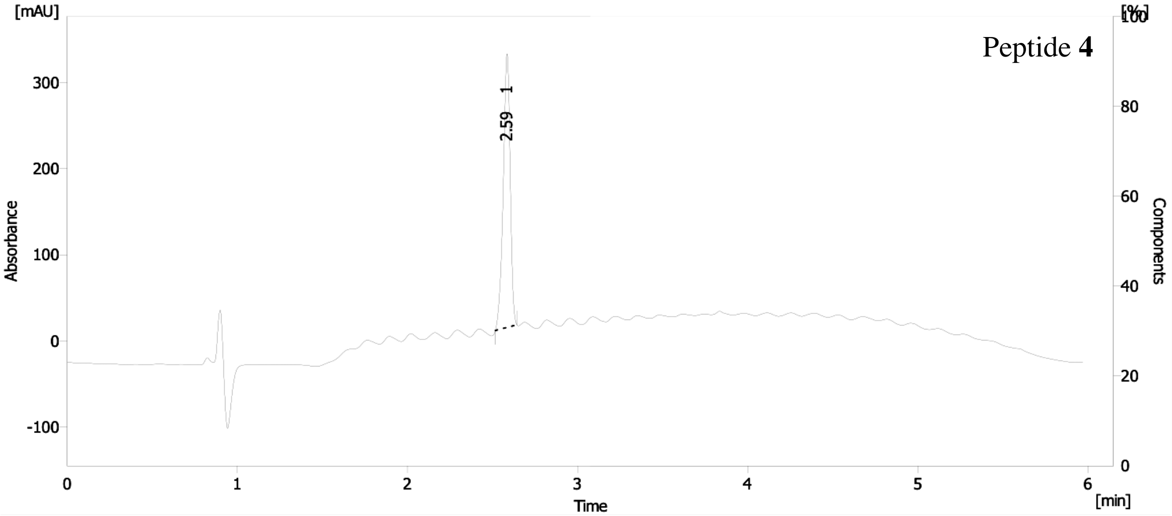
Peptide 4
[mAU]
[%]
Absorbance
Components
2.59 1
Time
[min]

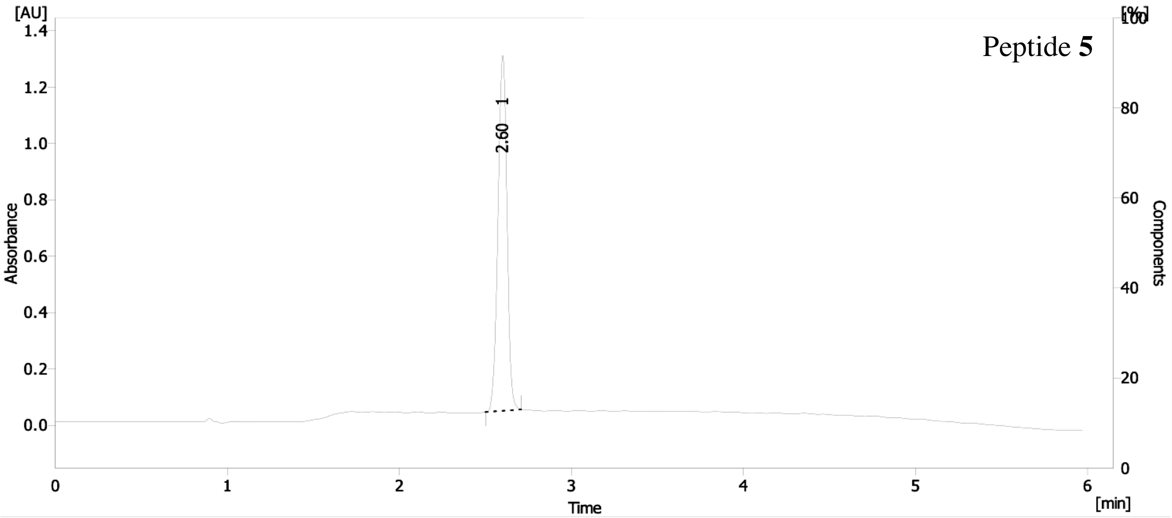
Peptide 5
[AU]
[%]
Absorbance
Components
2.60 1
Time
[min]

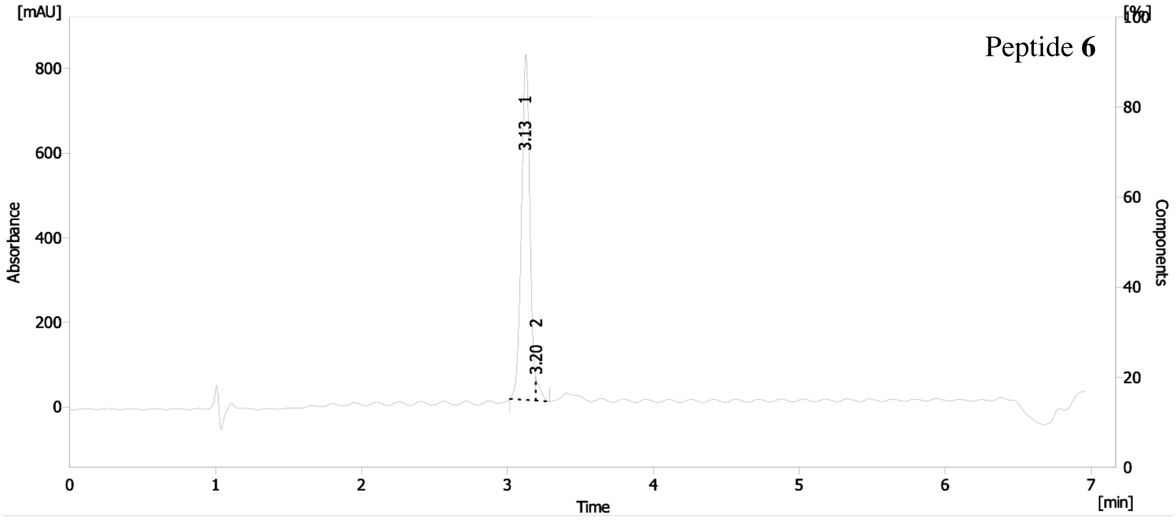

[mAU]
[%]
Peptide 6
800
600
400
200
0
Absorbance
3.13 1
3.20 2
100
80
60
40
20
0
Components
0
1
2
3
4
5
6
7
Time
[min]

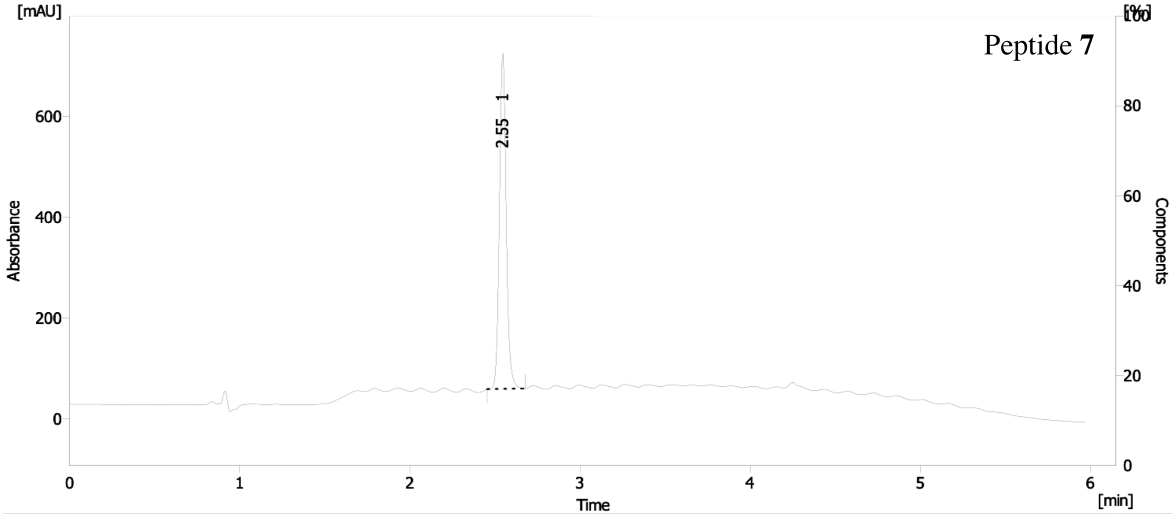

[mAU]
[%]
Peptide 7
600
400
200
0
Absorbance
2.55 1
100
80
60
40
20
0
Components
0
1
2
3
4
5
6
Time
[min]

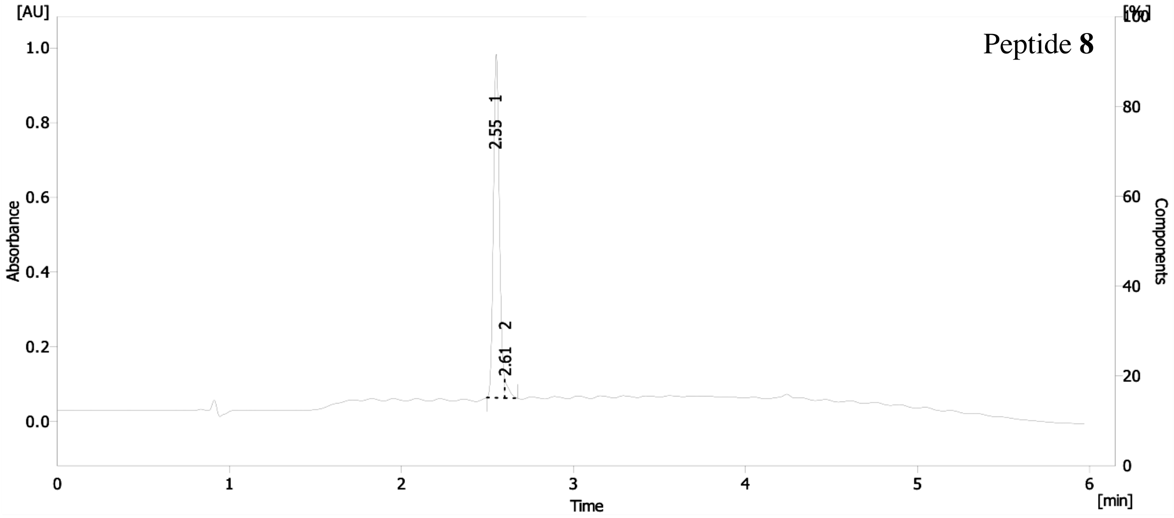

[AU]
[%]
Peptide 8
1.0
0.8
0.6
0.4
0.2
0.0
Absorbance
2.55 1
2.61 2
100
80
60
40
20
0
Components
0
1
2
3
4
5
6
Time
[min]

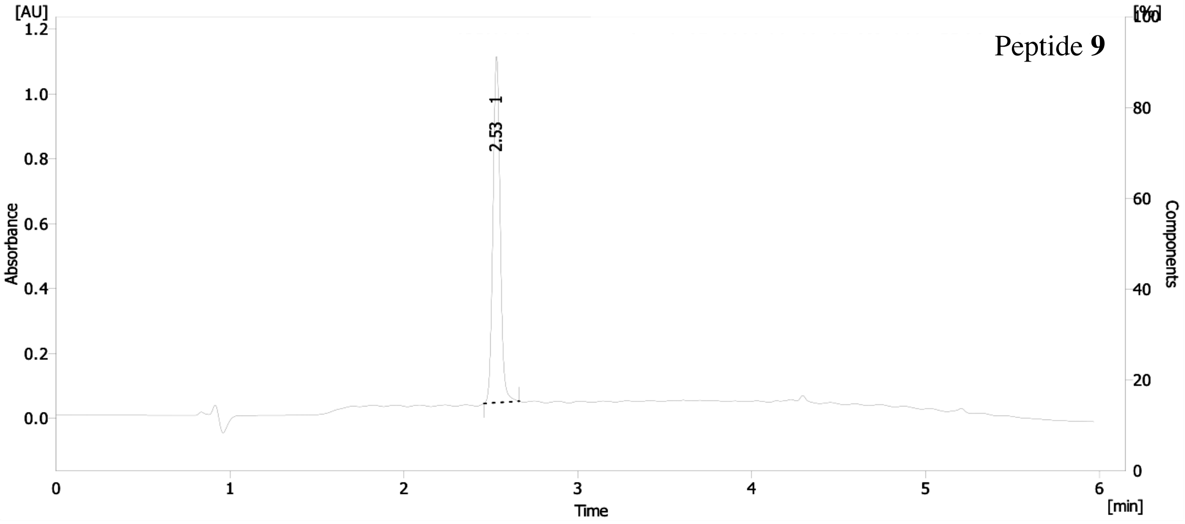
Peptide **9**
[AU]
Absorbance
Time
[min]
[%]
Components
2.53 1

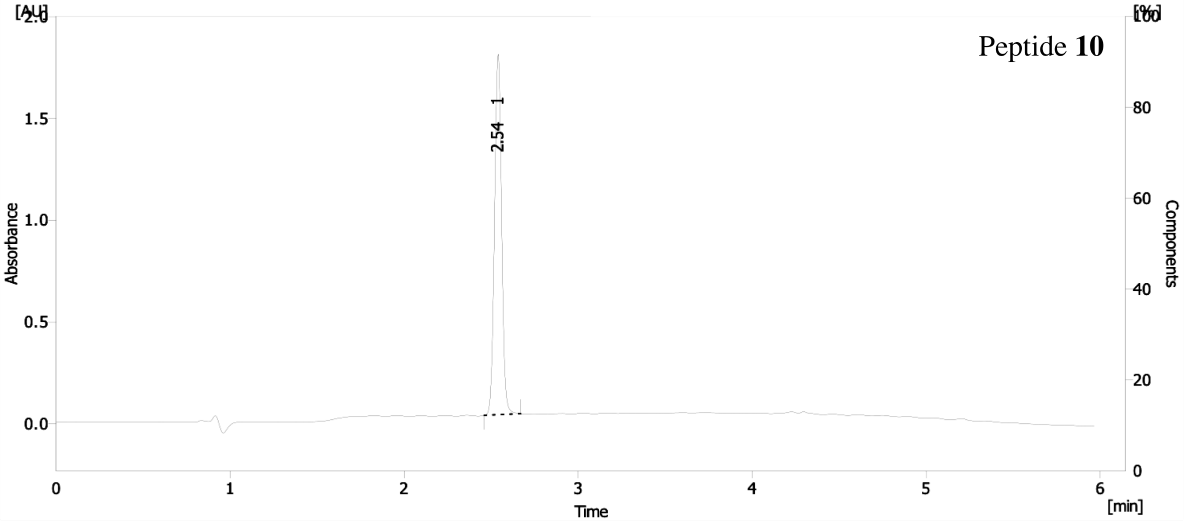
Peptide **10**
[AU]
Absorbance
Time
[min]
[%]
Components
2.54 1

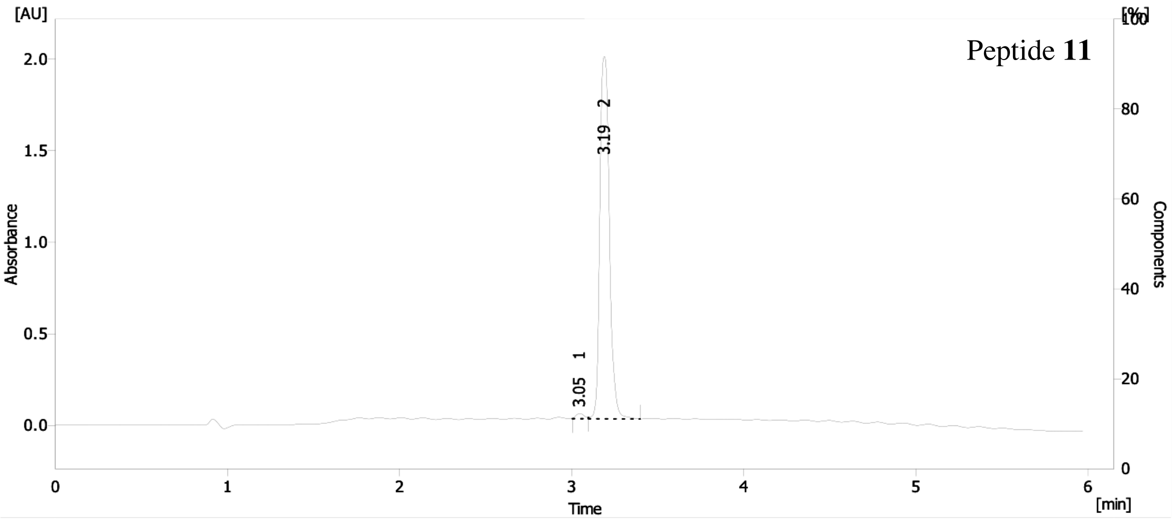
Peptide **11**
[AU]
Absorbance
Time
[min]
[%]
Components
3.05 1
3.19 2

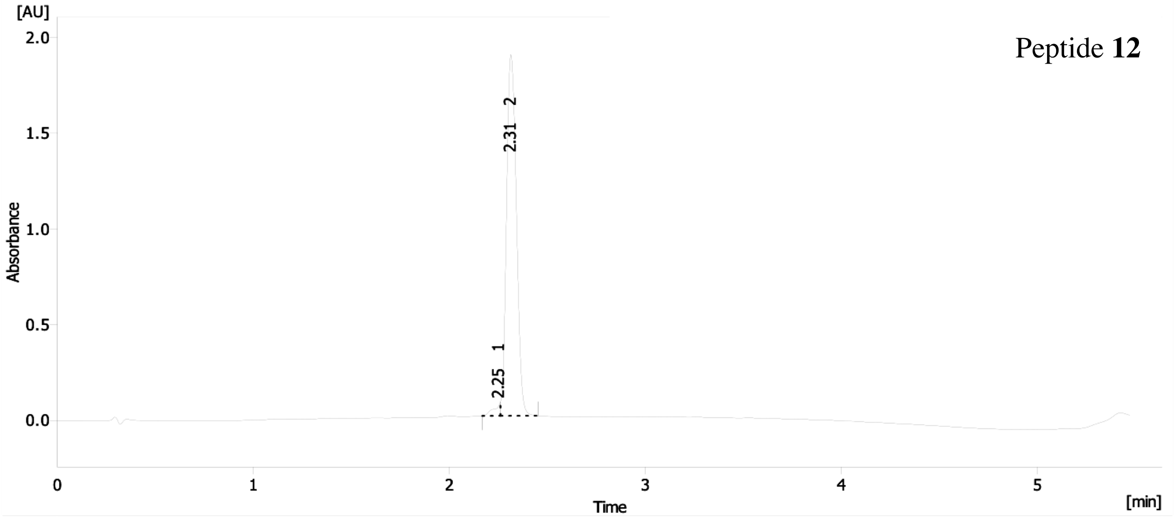

Peptide **12**

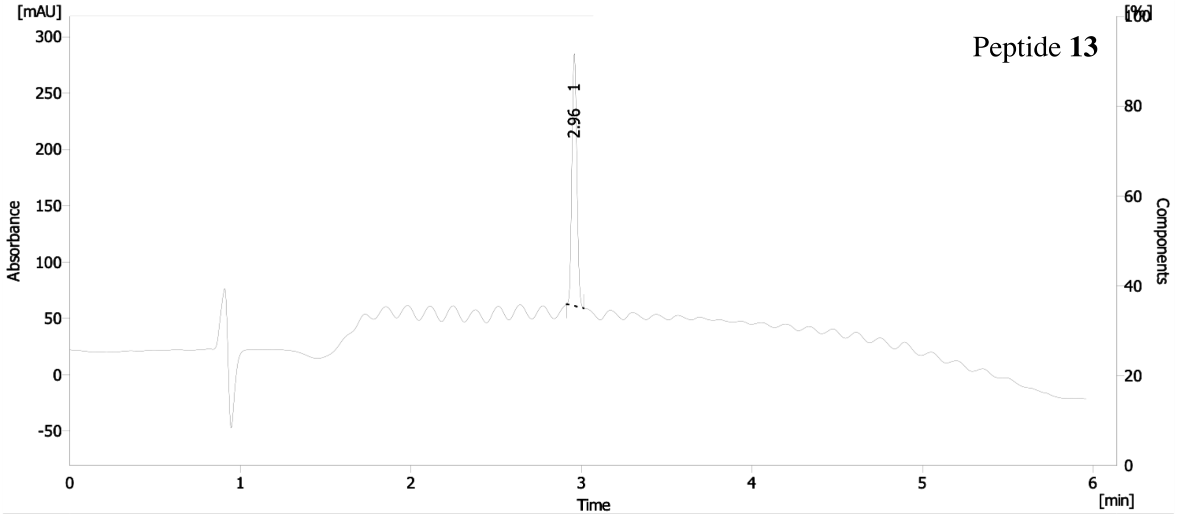

Peptide **13**

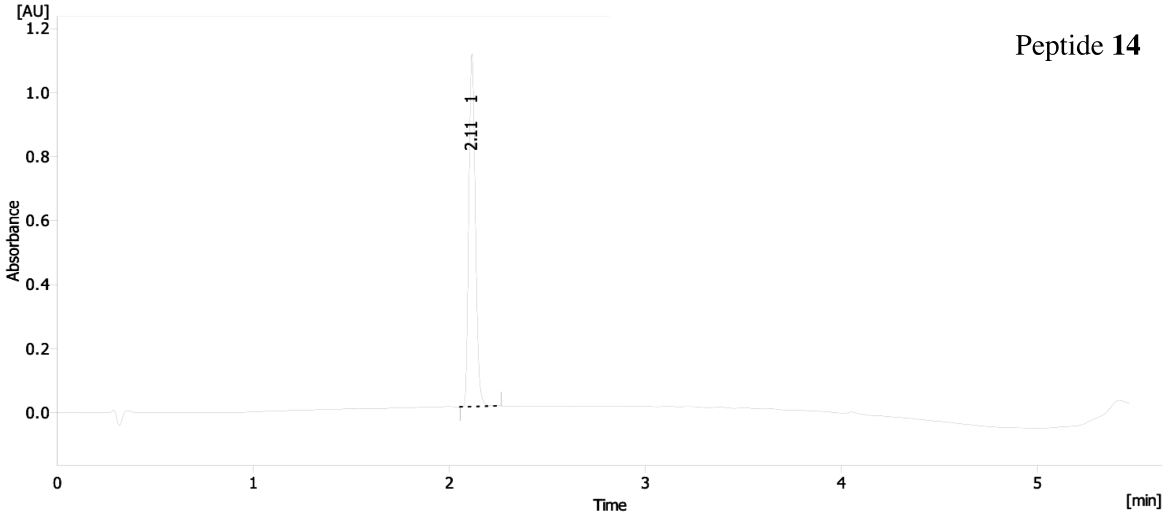

Peptide **14**

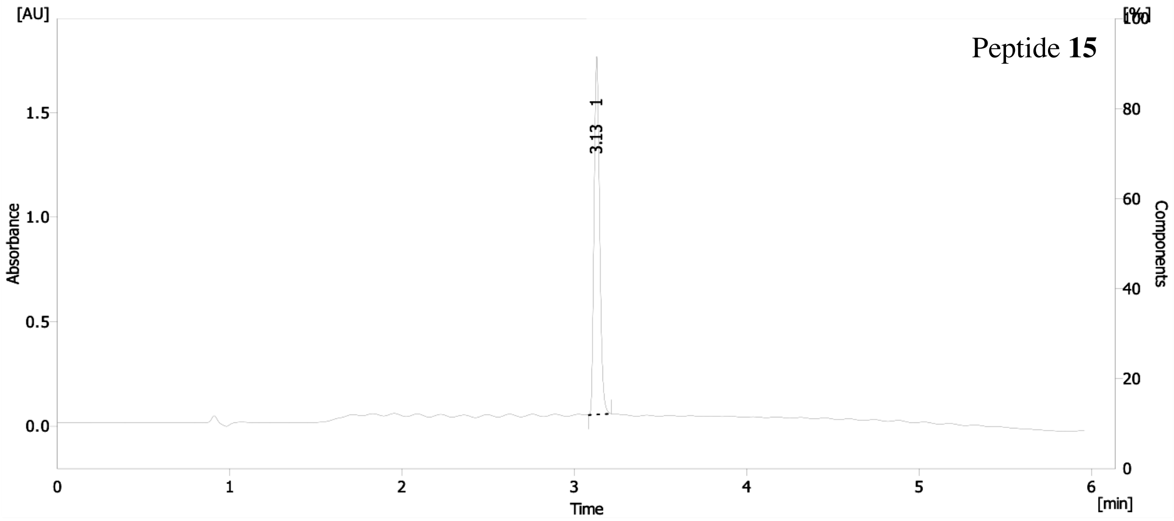
[AU]
Peptide 15
Absorbance
Components
[%]
3.13 1
Time
[min]

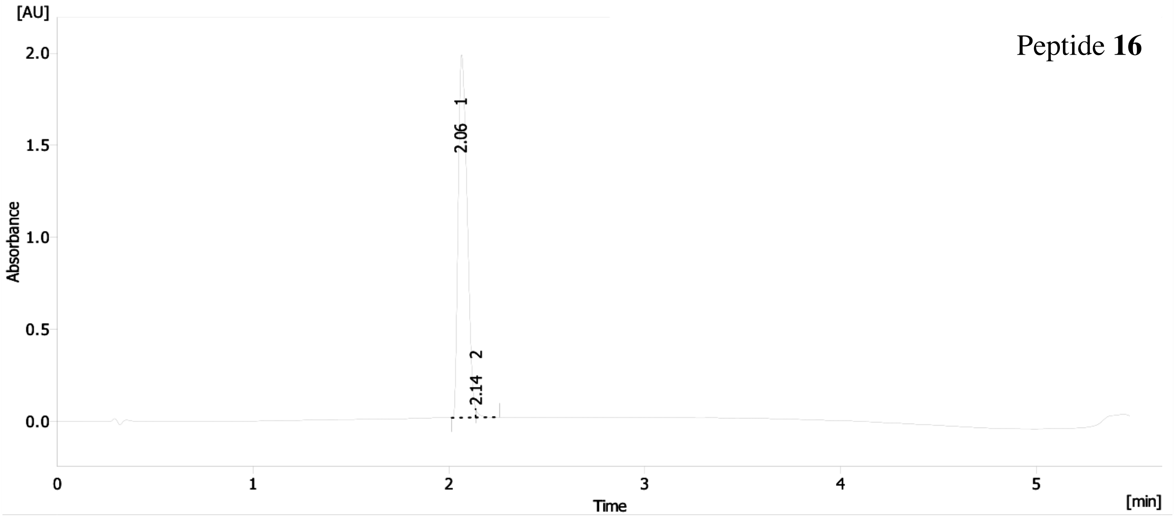
[AU]
Peptide 16
Absorbance
2.06 1
2.14 2
Time
[min]

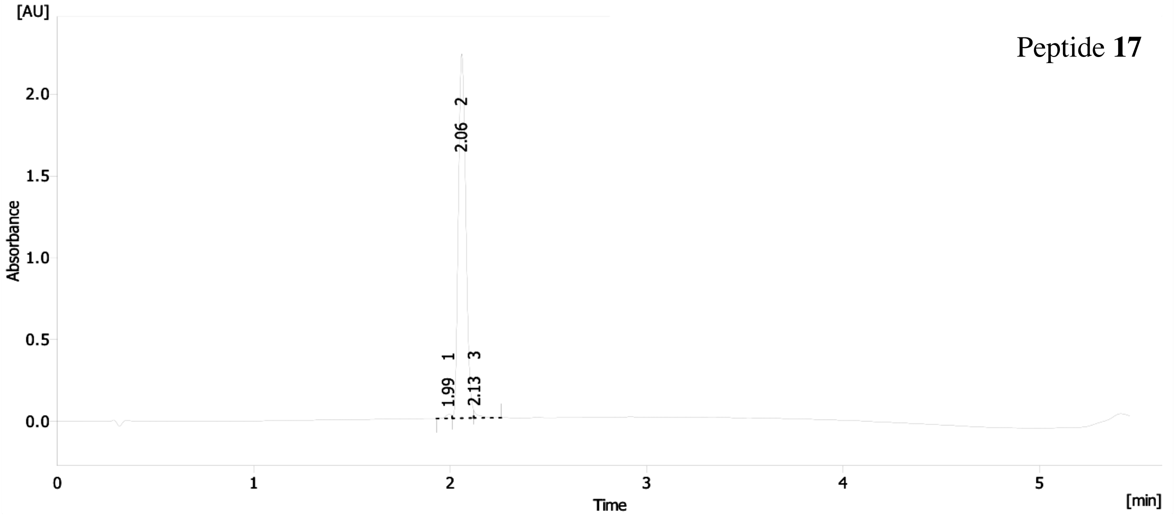
[AU]
Peptide 17
Absorbance
2.06 2
1.99 1
2.13 3
Time
[min]

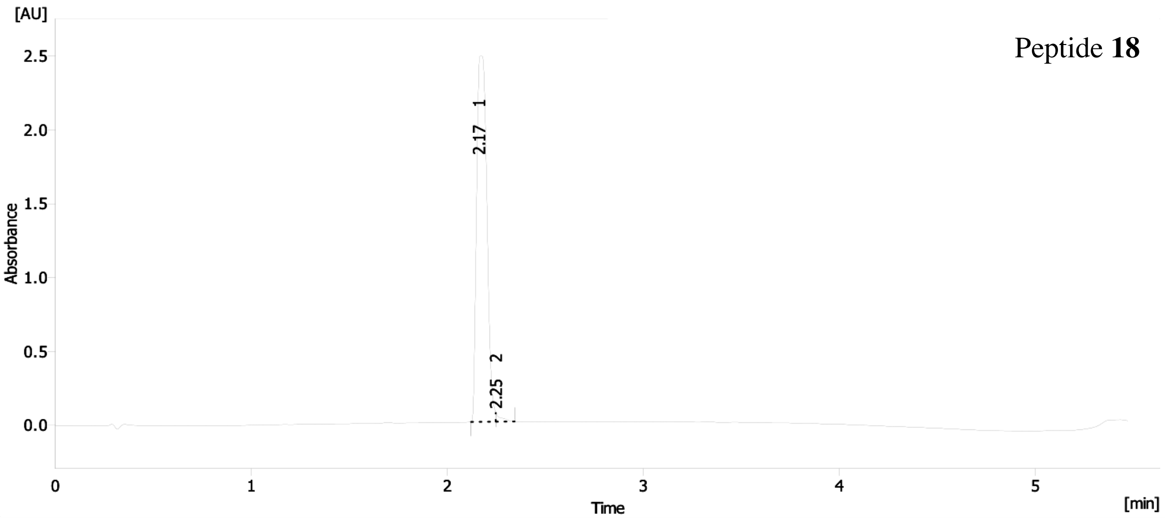

Peptide **18**

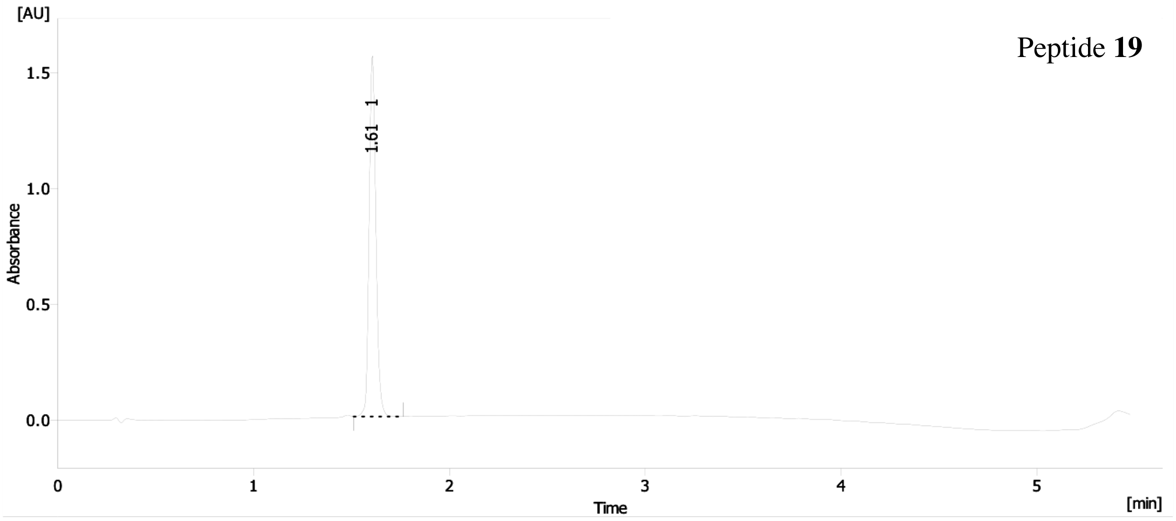

Peptide **19**

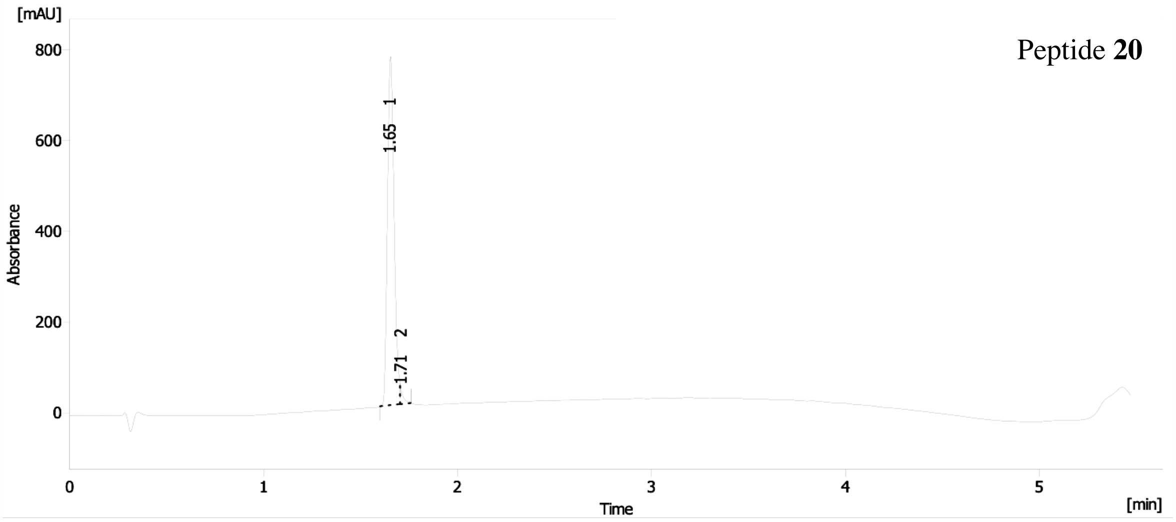

Peptide **20**

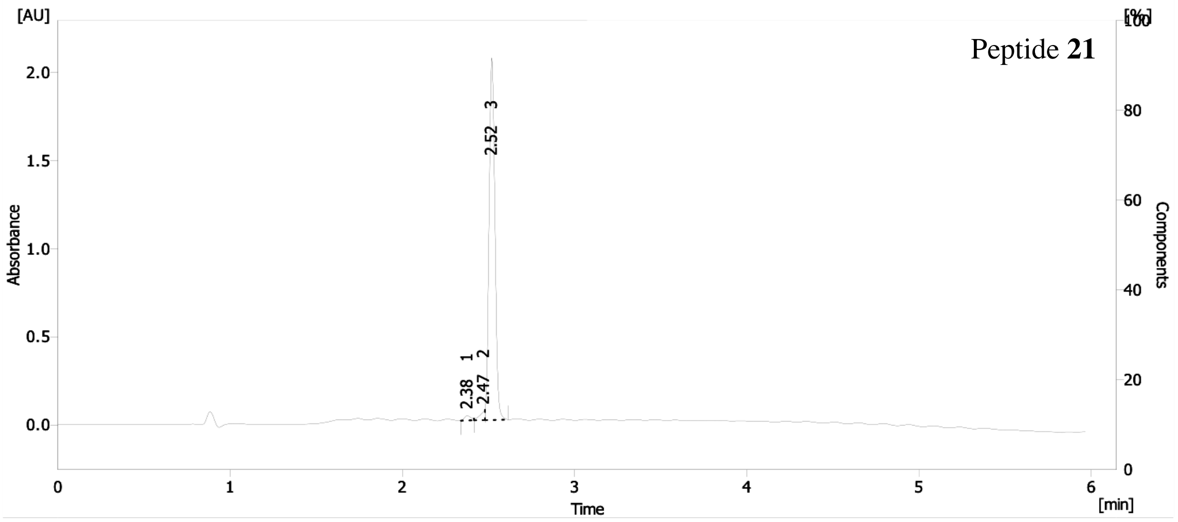
[AU]
[%]
Peptide **21**
2.52 - 3
2.38 - 1
2.47 - 2
Absorbance
Components
Time
[min]

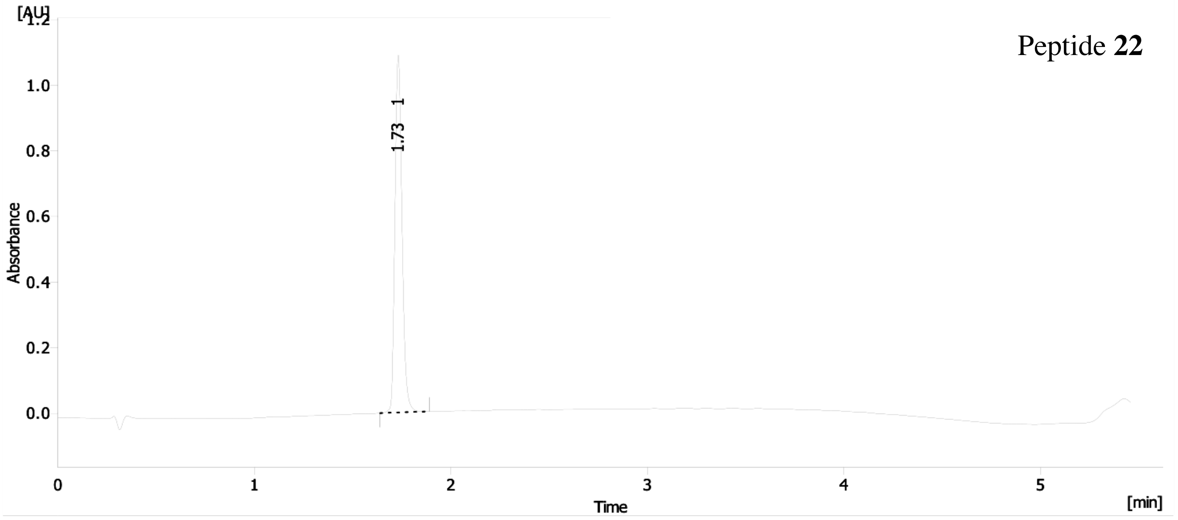
[AU]
Peptide **22**
1.73 - 1
Absorbance
Time
[min]

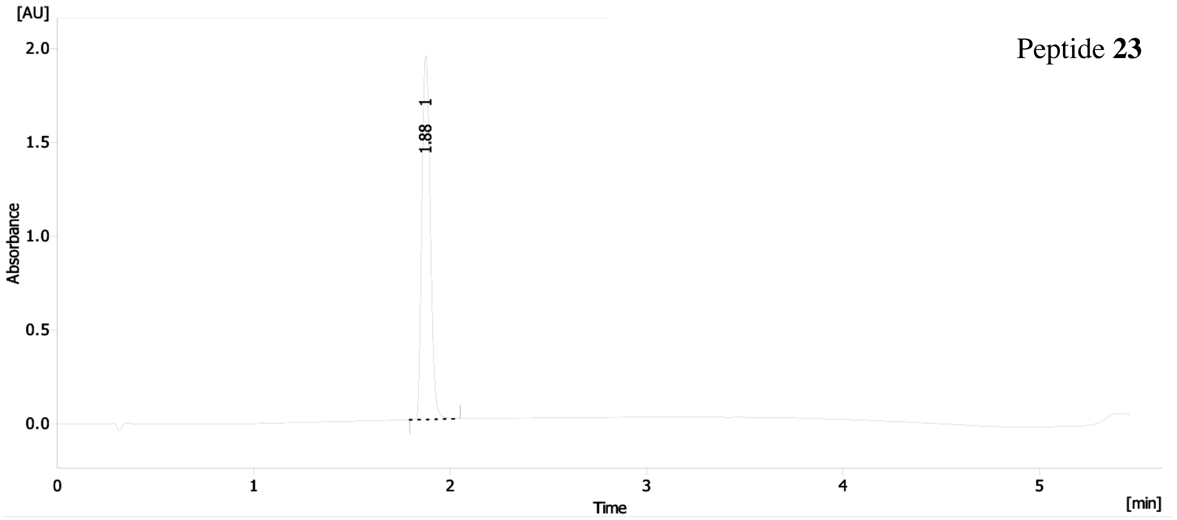
[AU]
Peptide **23**
1.88 - 1
Absorbance
Time
[min]

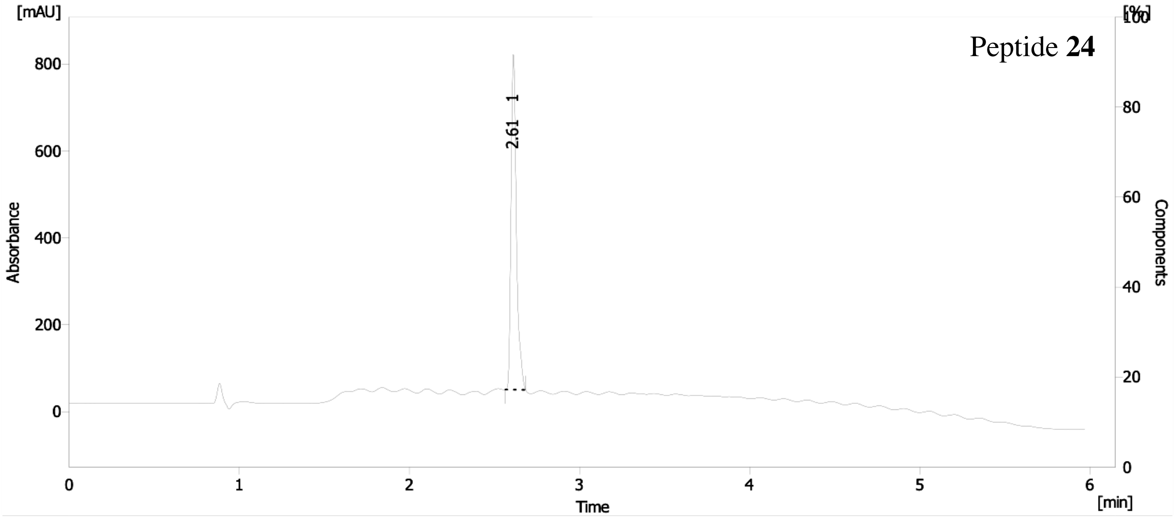
Peptide 24
[mAU]
[%]
100
800
600
400
200
0
80
60
40
20
Absorbance
Components
2.61 1
0
1
2
3
4
5
6
Time
[min]

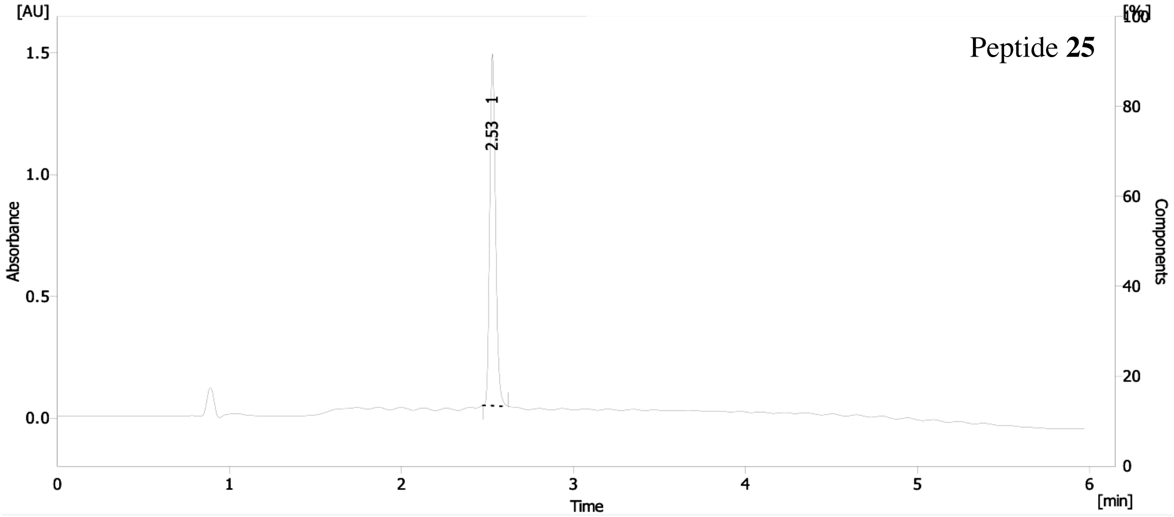
Peptide 25
[AU]
[%]
100
1.5
1.0
0.5
0.0
80
60
40
20
Absorbance
Components
2.53 1
0
1
2
3
4
5
6
Time
[min]

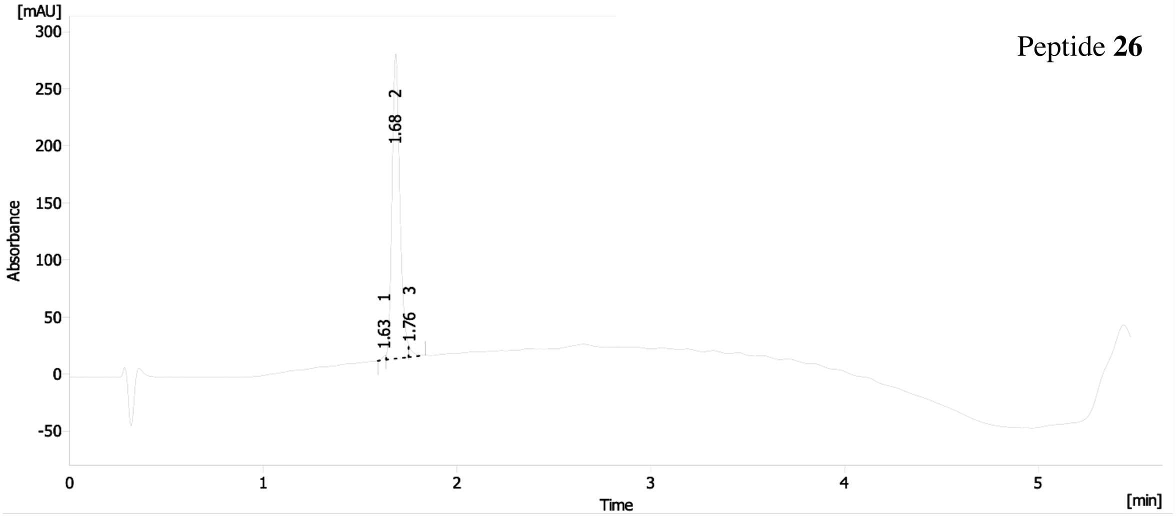
Peptide 26
[mAU]
300
250
200
150
100
50
0
-50
Absorbance
1.68 2
1.63 1
1.76 3
0
1
2
3
4
5
Time
[min]

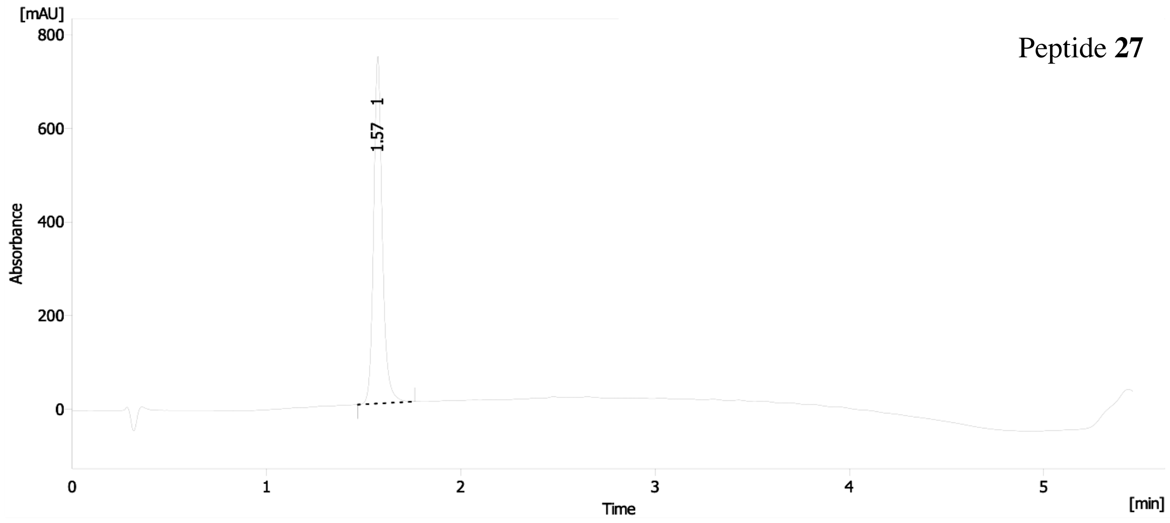
[mAU]
800
600
400
200
0
Absorbance
1.57 1
0
1
2
3
4
5
Time
[min]
Peptide 27

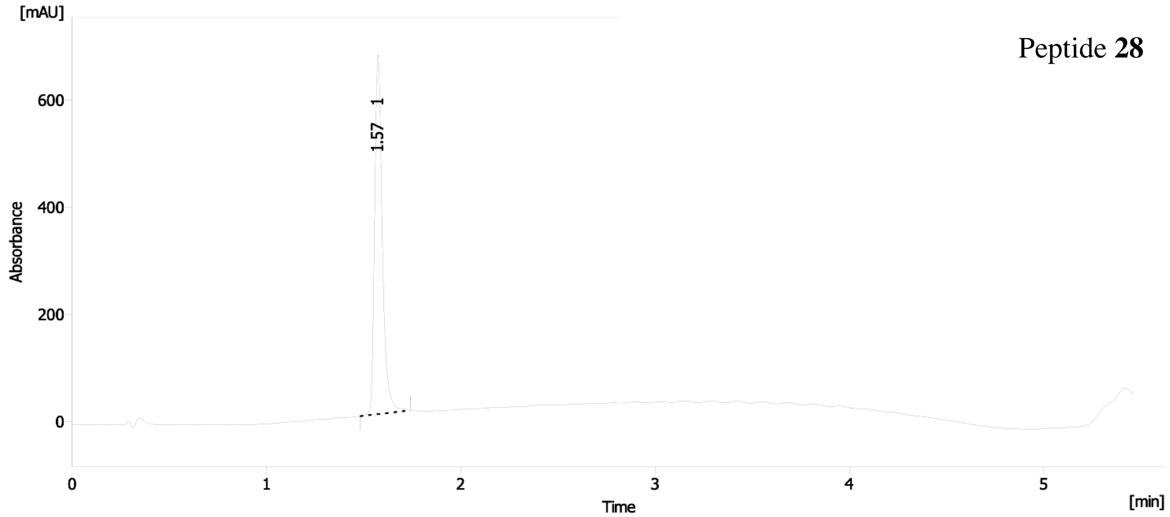
[mAU]
600
400
200
0
Absorbance
1.57 1
0
1
2
3
4
5
Time
[min]
Peptide 28

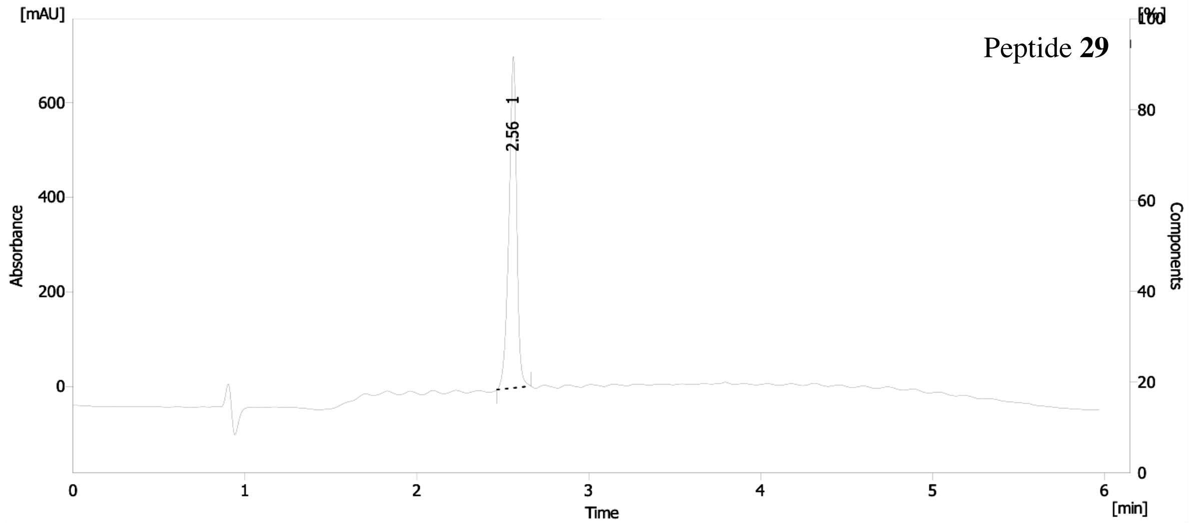
[mAU]
600
400
200
0
Absorbance
2.56 1
0
1
2
3
4
5
6
Time
[min]
[%]
100
80
60
40
20
0
Components
Peptide 29

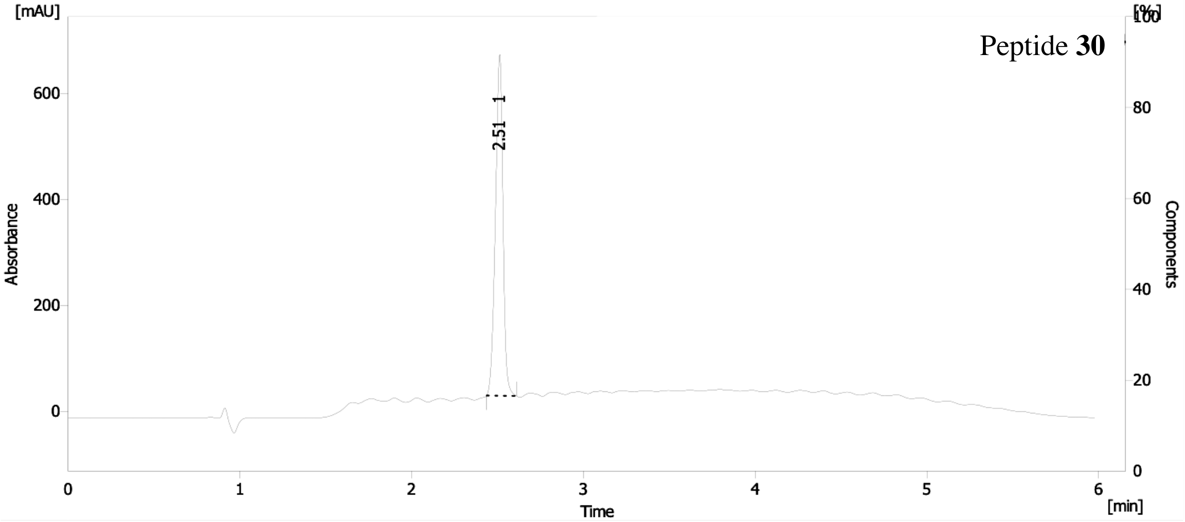
[mAU]
[%]
100
Peptide **30**
600
400
200
0
Absorbance
2.51 1
80
60
40
20
0
Components
0
1
2
3
4
5
6
Time
[min]

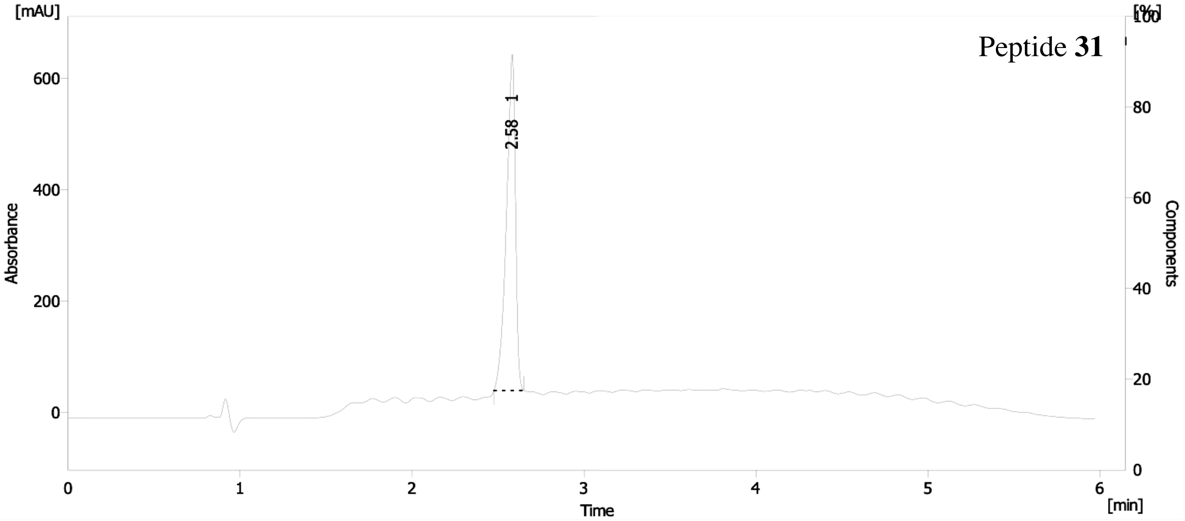
[mAU]
[%]
100
Peptide **31**
600
400
200
0
Absorbance
2.58 1
80
60
40
20
0
Components
0
1
2
3
4
5
6
Time
[min]

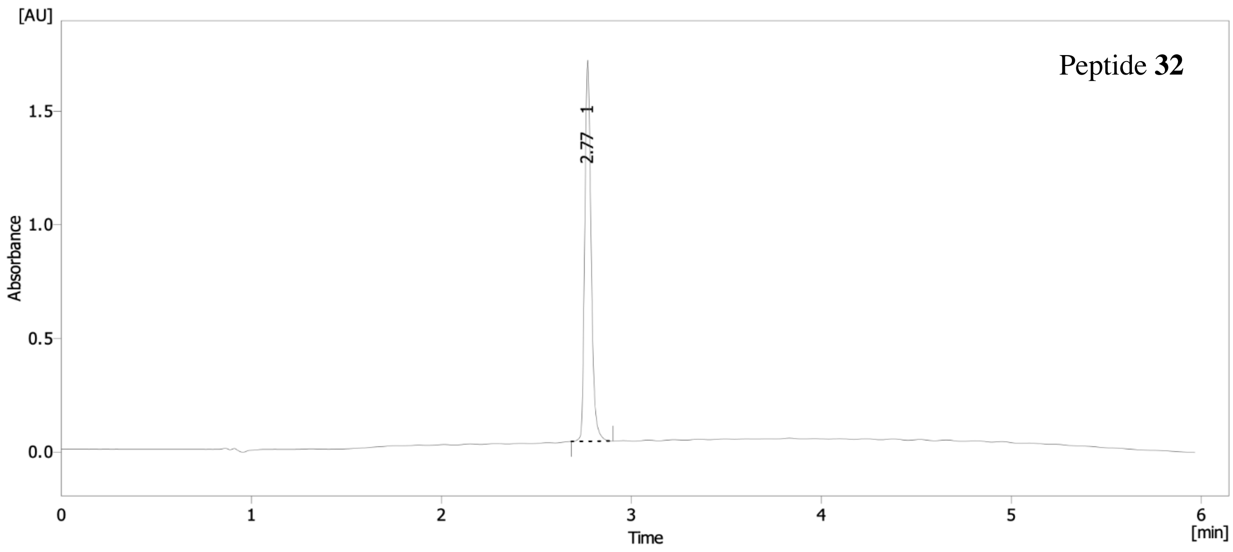
[AU]
Peptide 32
2.77 1
Absorbance
Time
[min]

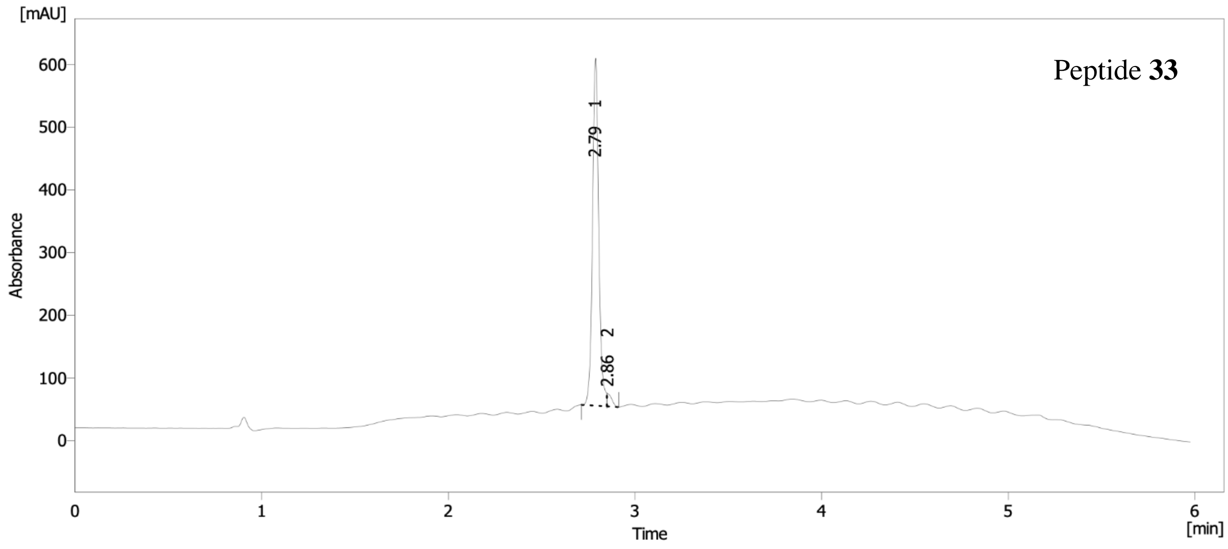
[mAU]
Peptide 33
2.79 1
2.86 2
Absorbance
Time
[min]

### 3 Protein production and purification

Codon-optimized synthetic genes encoding human HRAS (residues 1–169) and human SOS1 (residues 564–1049) were cloned as NdeI and XhoI fragments into pET28b (Novagen). His-tagged proteins RAS and SOS1 were expressed in *E. coli* strain BL21(DE3) and purified by immobilized metal ion affinity chromatography (IMAC) and size-exclusion chromatography as described previously by Boriack-Sjodin *et al.*,[3] and were conserved in a buffer containing 25 mM Hepes (pH 7.5), 150 mM NaCl and 10 mM $MgCl_2$.

RAS loaded with mant-GDP (mGDP) was obtained by mixing RAS with a 10-fold stochiometric excess of mGDP (Jena Bioscience) in a buffer containing an excess of ethylenediamine tetraacetic acid (EDTA, 20 mM). After overnight incubation at 4°C, an excess of $MgCl_2$ (50 mM) was added for 30 min and the free nucleotides were removed by gel filtration.

The RAS:SOS1 complex was formed by incubating RAS and SOS1 at a stochiometric ratio of 3:1 overnight at 4 °C with 20 mM EDTA and alkaline phosphatase (Roche). The complex was then purified on gel filtration.

### 4 Protein crystallization and Structure Determination

#### 4.1 Protein complexes crystallization and data collection

Crystals of RAS:SOS1 were obtained by the hanging-drop vapor-diffusion method at 277 K in 3.0 M sodium formate and 100 mM Tris buffer (pH 8.0) as described by Boriack-Sjodin *et al.*[3] Before being flash frozen in liquid nitrogen, all crystals were soaked in mother liquor supplemented with 20% glycerol and 10% peptide dissolved in DMSO.

1. RAS:SOS1:**2** was obtained by soaking with **2** at 10mM for 1 h. Data were collected at 100 K on the I24 beamline at the Diamond Light Source synchrotron (Oxfordshire, UK) with a wavelength of 0.97 Å and the structure was refined to 2.90 Å resolution.

2. RAS:SOS1:**3** was obtained by soaking with **3** at 5mM for 1 h. Data were collected at 100 K on the Proxima 1 beamline of the Soleil synchrotron (Saint-Aubin, France) with a wavelength of 0.98 Å and the structure was refined to 3.00 Å resolution.

3. RAS:SOS1:**4** was obtained by soaking with **4** at 5mM for 1 h. Data were collected at 100 K on the Proxima 2 beamline of the Soleil synchrotron (Saint-Aubin, France) with a wavelength of 0.98 Å and the structure was refined to 2.40 Å resolution.

4. RAS:SOS1:**5** was obtained by soaking with **5** at 5mM for 1 h. Data were collected at 100 K on the Proxima 2 beamline of the Soleil synchrotron (Saint-Aubin, France) with a wavelength of 0.98 Å and the structure was refined to 2.51 Å resolution.

5. RAS:SOS1:**10** was obtained by soaking with **10** at 5mM for 2 h. Data were collected at 100 K on the Proxima 2 beamline of the Soleil synchrotron (Saint-Aubin, France) with a wavelength of 0.98 Å and the structure was refined to 2.47 Å resolution.

##### 4.2 Structure determination and analysis

All diffraction data were integrated and scaled with XDS.[4] Models were built by iterative cycles of refinement with Phenix,[5] and manual building in Coot.[6] MolProbity was used for structure validation.[7] Data collection and refinement statistics are summarized in Table S2. The final Ramachandran statistics (% Favored:% Outlier) were 95.5:0.2, 93.0:0.7, 98.0:0.0, 97.3:0.0 and 97.7:0.0 for RAS:SOS1:**2**, RAS:SOS1:**3**, RAS:SOS1:**4**, RAS:SOS1:**5**, and RAS:SOS1:**10,** respectively. All structural figures were generated using PyMOL Molecular Graphics System, Version 2.3 Schrödinger, LLC (https://pymol.org/).

##### 4.3 Accession codes

PDB: X-ray coordinates for RAS:SOS1:**2** (8BE6), RAS:SOS1:**3** (8BE7), RAS:SOS1:**4** (8BE8), RAS:SOS1:**5** (8BE9), and RAS:SOS1:**10** (8BEA).

#### 5 Nucleotide exchange assays

Nucleotide exchange assays were conducted by measuring mGDP fluorescence (440 nm) on a *Tecan plate reader*. The reaction was started by mixing mGDP-loaded RAS (1 $\mu$M) with SOS (concentration varying from 0.1 to 1 $\mu$M), with a fixed volume (1/40) of peptide (concentrations varying from 0.24 to 2500 $\mu$M) and an excess of GTP (200 $\mu$M). Raw fluorescence data were fitted to a single exponential decay function, and activation rates were obtained relative to the derived rate obtained with RAS:SOS1 alone. The maximal activation and $EC_{50}$ values were calculated by plotting derived rates as a function of ligand concentration and fitting using a four-parameter dose-response curve using Prism 8 (Graph pad Software Inc.).

#### 6 Biolayer Interferometry

Biolayer interferometry measurements were performed using Streptavidin biosensors on an Octet Red96 (Forte Bio, Inc.) system at 25 °C, shaking at 1000 rpm and in a buffer containing 25mM Hepes pH 7.5, 150 mM NaCl, 10 mM $MgCl_2$, 0.1% BSA and 0.005% Tween 20. Following a first step of equilibration with buffer, the biosensors were loaded with biotinylated peptide at a concentration of 10 $\mu$g/mL to reach an amplitude signal of 1. After baseline acquisition, the biosensors were transferred to wells supplemented with increasing concentrations of SOS1 (300 to 0.14 $\mu$M with a 1:3 serial dilution) to measure the association step. Finally, the biosensors were transferred to wells containing buffer to measure the dissociation step. All samples were prepared and measured in duplicate. To obtain an apparent dissociation constant, the amplitude signal at equilibrium was normalized and was plotted vs. the protein concentration. The resulting titration curve was fitted on a Langmuir equation.

#### 7 Kinetics

$K_1$ and $k_2$ values were determined by measuring mGDP (Jena Bioscience) fluorescence (excitation at 350 nm, emission filtered at 400 nm) on a stopped-flow apparatus (Applied Photophysics). The reaction was started by mixing mGDP loaded RAS (0.5 $\mu$M) with SOS1 (varying concentrations from 0 to 50 $\mu$M), an excess of peptide (500 $\mu$M), and an excess of GDP (100 $\mu$M). Raw fluorescence data were fitted to a single exponential decay function. $K_1$ and $k_2$ values were calculated by plotting derived rates as a function of SOS1 concentration with the equation indicated in Figure S7 using Prism 8 (Graph pad Software Inc.).

$K_2$ and $k_{-1}$ values were determined by measuring mGDP (Jena Bioscience) fluorescence (excitation at 350 nm, emission filtered at 400 nm) on a stopped-flow apparatus (Applied Photophysics). Reaction was started by mixing SOS1-RAS (0.5 $\mu$M) with an excess of peptide (500$\mu$M) and mGDP (varying concentrations from 0.5 to 25 $\mu$M). Raw fluorescence data were fitted to a single exponential association function. $K_2$ and $k_{-1}$ values were calculated by plotting derived rates as a function of mGDP concentration with the equation indicated in Figure S7 using Prism 8 (Graph pad Software Inc.).
